# Nanodomain distribution and function of PIN-FORMED auxin efflux carriers in the plasma membrane of tobacco cells are defined by their interactions with the cell wall

**DOI:** 10.1101/2025.01.23.634346

**Authors:** Ayoub Stelate, Kateřina Malínská, Eva Tihlaříková, Karel Müller, Roberta Vaculíková, Martina Knirsch, Zuzana Vondráková, Erik Vlčák, Katarzyna Retzer, Michaela Neubergerová, Roman Pleskot, Karel Harant, Vlada Filimonenko, Kateřina Schwarzerová, Vilém Neděla, Jan Petrášek

## Abstract

Plant development is under the morphogenic control of auxin. In addition to biosynthesis and metabolism, auxin concentration gradients are maintained by directional intercellular transport via PIN-FORMED (PIN) auxin efflux carriers. Although the structure-function properties of PINs have been described, it is still unclear whether individual members of PIN family could localize differently within the plasma membrane (PM) and whether their nanodomain distribution defines their function. To address this, we used cultured tobacco cells (*Nicotiana tabacum* L., cv. BY-2) and revealed by RNA-seq and RT-qPCR the cell stage-specific presence of transcripts of tobacco PIN homologs. We used high-resolution light microscopy and two independent immunoelectron microscopy techniques in cell lines expressing functional GFP-tagged inducible versions of *Nt*PIN11T, *Nt*PIN2T and *Nt*PIN3bT. We show that *Nt*PINs are distributed within specific PM nanodomains and that removal of the cell wall alters their appearance. By a series of *in vivo* microscopic observations, we provide evidence that *Nt*PIN11T is the most homogenously distributed and the least mobile *Nt*PIN. Pharmacological treatments suggested that the immobilization of *Nt*PIN11T depends on the actin and microtubular cytoskeleton and the cell wall composition. Using comparative co-immunoprecipitation (co-IP) analysis of isolated membrane fractions for all three *Nt*PINs we finally identified several novel interaction partners indicating a preferential association of *Nt*PIN11T with cell wall GPI-anchored arabinogalactan proteins. In conclusion, our results suggest a model in which specific immobilization of PINs through interactions with the cell wall affects their function.

## Introduction

Numerous functions of plant hormone auxin, represented mainly by indole-3-acetic acid (IAA), are connected with its transmembrane transport within plant tissues (Friml, 2022). Although IAA, depending on its dissociation rates, could be transported across membranes by diffusion to generate developmentally instructive auxin maxima, specific IAA uptake carriers from AUX1/LAX family cooperate with IAA efflux carriers from ABCB and PIN-FORMED (PIN) families (Band et al., 2014; Kramer et al., 2011; Mellor et al., 2022) PIN proteins are typically known for their asymmetric localizations within PM, which is very important in the IAA directional transport (Adamowski & Friml, 2015; Marhava, 2022). Their localization and function depend on regulatory phosphorylations (Bassukas et al., 2022), which help these proteins to react very rapidly on IAA (Dubey et al., 2021). The crossover elevator-type mechanism of PIN-mediated transport shared with archaeal organisms (Su et al., 2022; Ung et al., 2022; Yang et al., 2022) was identified recently, providing a long time awaited evidence on the mechanism of their action and regulation of this mechanism.

PINs display tissue-specific distributions within their gene expression domains (Vanneste & Friml, 2009; Vieten et al., 2005). The polarity of cellular PIN localization also reflects sequence-specific signals within individual PIN proteins (Wiśniewska et al., 2006). The distribution of PINs within PM is not homogeneous (Kleine-Vehn et al., 2011) and they are also highly dynamic through clathrin-mediated endocytosis (Kitakura et al., 2011; Narasimhan et al., 2020). Fluorescence microscopy techniques (Jacobson et al., 2019) showed that plant PM consists of a patchwork of many subdomains that coexist on a large temporal and spatial scale (Bücherl et al., 2017; Platre et al., 2019). PM nanodomains have been implicated in the regulation of endocytosis in reaction to external stimuli and clustering of proteins, and they could also regulate the function of carriers (Martinière & Zelazny, 2021). Since root and shoot tissues of *Arabidopsis thaliana* are not well accessible for demanding advanced microscopy methods that provide both high spatial and temporal resolution (Platre et al., 2019), there are still quite a few studies describing how cellular structures in close contact with plant PM define function and localization of PINs (Li et al., 2021).

In this work, we used several molecular biology and microscopy techniques in the model of cultured tobacco BY-2 cells (Nagata et al., 1992). This allowed us to compare individual tobacco PIN proteins in the same cellular context of highly homogeneous plant cell populations. We show that GFP-tagged inducible versions of all tested tobacco auxin efflux carriers transport auxin, they all are present in PM nanodomains, and the removal of the cell wall changes their appearance within PM. *Nt*PIN11T as the most homogenously distributed is shown to be the least mobile by a set of *in vivo* microscopic observations. This immobilization depends on intact cytoskeleton and cell wall composition, which seems to regulate the function of this carrier. We support these results by the identification of several novel interactors that have been identified by co-IP analysis of solubilized PM fractions. Our results suggest that within the canonical group of PINs the function of individual members is influenced by their specific cytosolic, PM, and cell wall interactions.

## Results and discussion

### *Nt*PIN11T is the main canonical PIN in tobacco BY-2 cells

Our previous results clearly showed that genes for PIN auxin efflux carriers are transcribed in dividing tobacco BY-2 cells and their mRNA levels decrease upon auxin removal (Müller et al., 2019, 2021). Therefore, we first compared the transcription of all *NtPINs* in exponential and stationary cells by RNA-seq analysis in 2-day-old dividing cells and 7-day-old elongated cells. Transcripts of all four *NtPIN* genes were readily detectable during both growth phases, with *NtPIN11*, a member of a sister of *PIN1* (*SoPIN1*) group (O’Connor et al., 2017), being the most abundant (Fig. 1*A*). We supported these results by RT-qPCR transcriptional profiling of *NtPIN*s during the life cycle (*SI Appendix*, Fig. S1*A*), where the *NtPIN11* transcription exhibited remarkable stability over the course of 10 days of cultivation. In contrast, *NtPIN2* transcription increased transiently during the exponential growth phase and decreased in the stationary phase. Both *NtPIN3a* and *NtPIN3b* were transcribed slightly more in elongated cells. To understand if these life cycle stage-specific transcriptions of individual *NtPINs* are also reflected on the level of the cell cycle, we performed RT-qPCR analysis in synchronized cell populations (*SI Appendix*, Fig. S1*B*). Interestingly, for *NtPIN2* we detected a significant increase of the transcription timely correlating with the mitotic peak and M/G1 transition, while the transcription of all other *NtPIN* genes increased during G1. Early cytokinetic and post-cytokinetic functions of PIN2 were previously shown in *Arabidopsis thaliana* to involve specific posttranslational interaction with protein kinases (Glanc et al., 2019) or dynamin GTPase DRP1a (Mravec et al., 2011). Our data thus support the idea that the level of the particular *PIN* gene expression is part of a compensatory mechanism, which was shown previously to regulate the transcription of individual *Arabidopsis thaliana PIN* genes (Vieten et al., 2005). The preferential presence of *NtPIN11* transcripts could also reflect the sub-functionalization of PIN1-type PINs, shown in monocots (O’Connor et al., 2017) and tomato (Martinez et al., 2016).

**Fig. 1.**
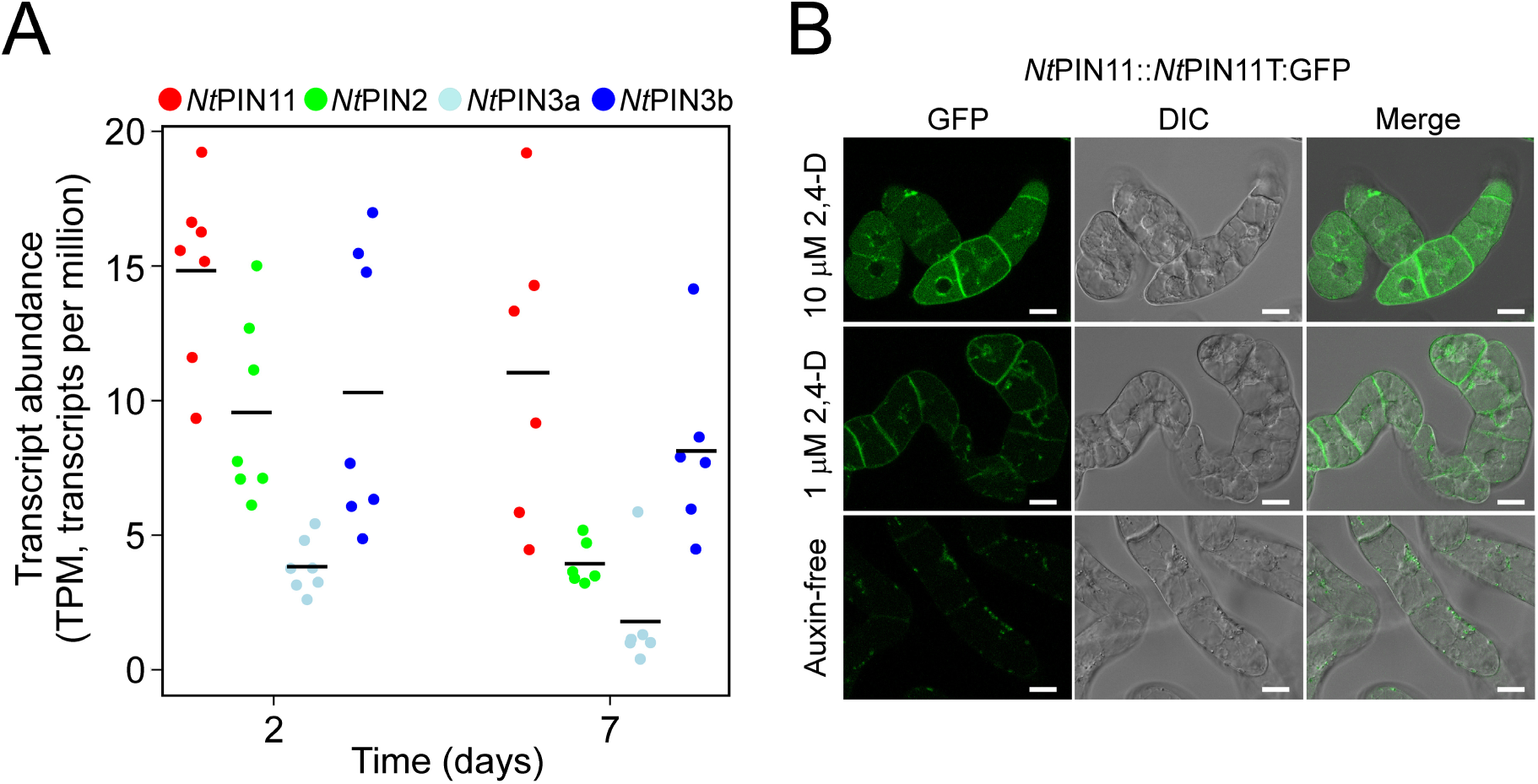
Expression analysis of individual *Nt*PINs in tobacco cells. (*A*) Transcript abundances for *NtPIN11*, *NtPIN2*, *NtPIN3a* and *NtPIN3b*, RNA-seq data from cells in exponential (2-day-old) and stationary (7-day-old) growth phase. Individual values represent the sum abundances of transcripts for both S (from parental *Nicotiana tomentosiformis*) and T (from parental *Nicotiana sylvestris*) forms of the particular *NtPIN* gene. The number of independent biological repetitions, i.e. independent RNA-seq analyses: 7 (day 2), 6 (day 7). Horizontal black lines indicate the mean. The color coding is indicated in the legend above the plot. (*B*) Localization of *Nt*PIN11T::*Nt*PIN11T:GFP in 2-day-old cells. Confocal and DIC microscopy. Middle row, cells cultured in control medium with 1 µM 2,4-D; upper row, cells cultured in medium with 10 µM 2,4-D; lower row, cells cultured in auxin-free medium.

To investigate whether the transcription of specific *NtPIN* gene is reflected on the protein level, we expressed GFP-tagged *NtPINs* (Müller et al., 2019) under native promoters and studied their intracellular distribution using confocal microscopy. We detected a GFP signal only for *Nt*PIN11T::*Nt*PIN11T:GFP (Fig. 1*B*, middle row), while there was no signal observed in any of the 40 tested transformed lines for *Nt*PIN2T::*Nt*PIN2T:GFP and *Nt*PIN3bT::*Nt*PIN3bT:GFP. Importantly, the fluorescence of *Nt*PIN11T::*Nt*PIN11T:GFP at the PM was stimulated after the addition of 10 µM 2,4-D (Fig. 1*B*, upper row), and completely disappeared in an auxin-free medium (Figure 1*B*, lower row), supporting our previously shown drop of *NtPINs* transcription in auxin-starved cells (Müller et al., 2019, 2021).

Altogether, although all three types of *NtPINs* were actively transcribed, we detected only *Nt*PIN11T on the protein level, suggesting possible differences in the stability of individual *Nt*PINs within PM. Therefore, we decided to analyze the localization and dynamics of individual *Nt*PINs in high spatial and time resolution.

### Removal of the cell wall decreases the homogeneity in the PM distribution of *Nt*PINs

To compare individual *Nt*PIN localization in the same cellular context, we used previously published cell lines carrying *Nt*PIN11T-GFP, *Nt*PIN2T-GFP, and *Nt*PIN3bT-GFP under the control of a β-estradiol receptor-based chemically-inducible system (Müller et al., 2019; Zuo et al., 2000). Upon induction, these lines displayed very clear PM localization of all three studied *Nt*PINs (Müller et al., 2019). Before investigating the dynamics and localization patterns of *Nt*PINs within the PM at the high spatial and temporal resolution, cells were induced with β-estradiol and auxin transport was assayed after 48 h. In all three lines, we observed a significant decrease in the accumulation of the auxin efflux carrier substrate [^3^H]NAA (Delbarre et al., 1996) to approximately half the level (Fig. 2*A*), indicating that the overexpressed *Nt*PINs are all functional auxin efflux carriers. Protein 3D structures generated by AlphaFold2 (Jumper et al., 2021; Mirdita et al., 2022) show that the GFP insertion in the cytosolic loop has a good level of freedom (*SI Appendix*, Fig. S2*A*). Furthermore, auxin transport activities of *Nt*PINs were tuned in a dose-dependent manner by increasing the concentration of β-estradiol, as shown for *Nt*PIN11T-GFP and XVE-*Nt*PIN2T-GFP (*SI Appendix*, Fig. S2*B*), and previously also for *Arabidopsis thaliana* PIN3 and PIN7 heterologously expressed in tobacco cells (Kashkan et al., 2022).

**Fig. 2.**
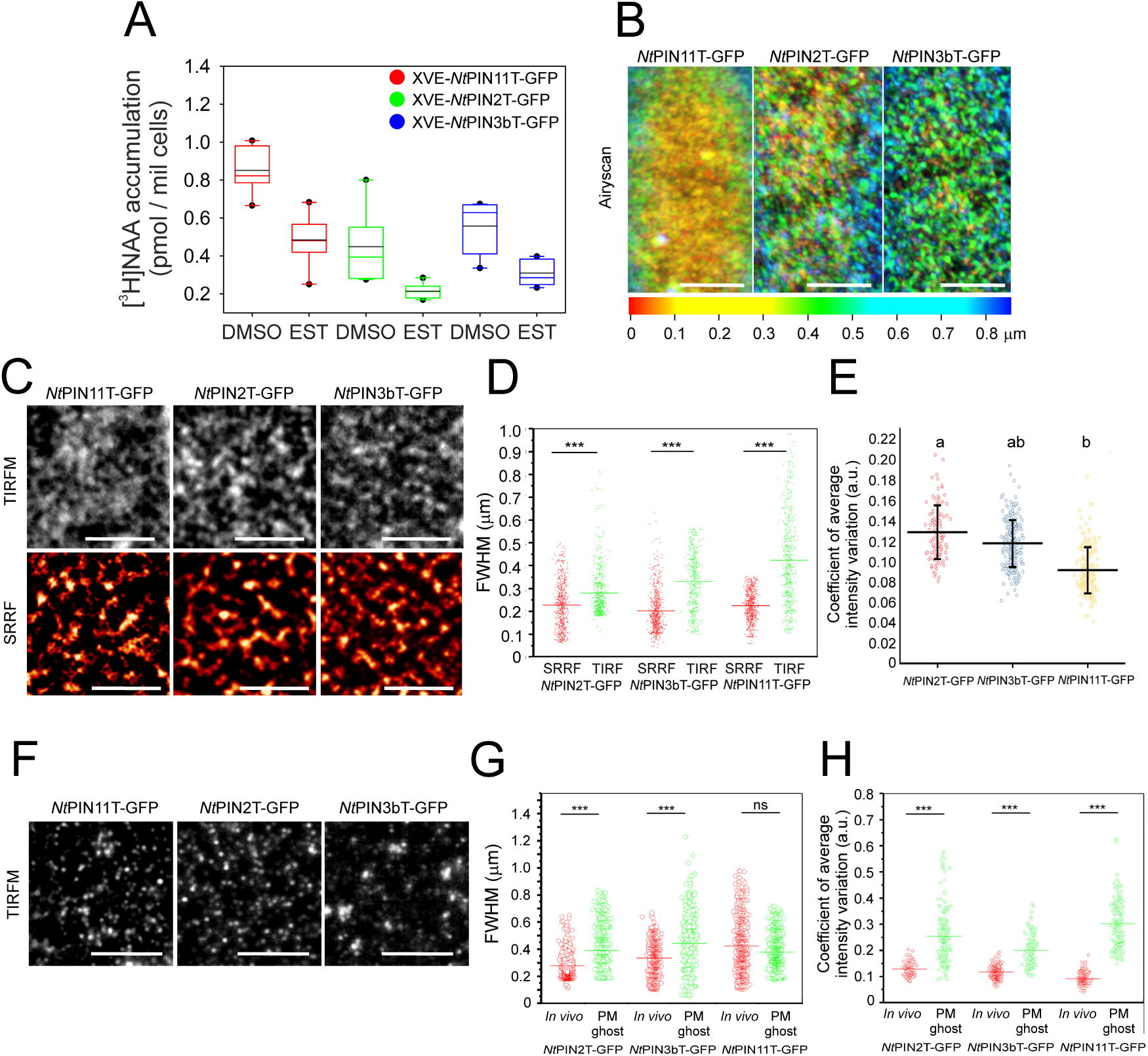
Functional analysis and PM distribution of GFP-tagged *Nt*PINs. (*A*) Functional analysis of inducible *Nt*PINs in XVE-*Nt*PIN11T-GFP, XVE-*Nt*PIN2T-GFP and XVE-*Nt*PIN3bT-GFP cell lines. Induction with 1 µM β-estradiol for 48 h (EST) significantly decreased the [^3^H]NAA accumulation in comparison with DMSO-treated cells (DMSO) for all tested *Nt*PINs. Data from six biological repetitions (independent inductions in the particular line) are shown with a mean (black lines) and median (color lines). Data points outside the 1.5 times interquartile range from 1st/3rd quartile are shown as outliers. In all studied *Nt*PINs, the significance at P < 0.05 was tested positively with Mann-Whitney U-test for induced versus non-induced cells. (*B*) Depth color-coded stacks from 5 Airyscan sections through the surface of tobacco cells (170 nm each) for three GFP-tagged *Nt*PINs. Note that nanodomains are not distributed homogenously along the z-axis, with only *Nt*PIN11T-GFP showing a packed appearance within a thin cortical region at the surface of the cells containing PM and associated structures. (*C*) *In vivo* TIRFM and SRRF images of GFP-tagged *Nt*PINs nanodomains. (*D*) Comparison of nanodomain dimensions imaged by TIRFM and SRRF (FWHMs). Data from 20 individual cells for each protein, n=582/643 (TIRFM/SRRF, *Nt*PIN11T), 580/623 (TIRFM/SRRF, *Nt*PIN2T), and 650/654 (TIRFM/SRRF, *Nt*PIN3bT). The significance of the difference in FWHM for TIRFM and SRRF is tested with T-test; horizontal lines show the mean values, ***P < 0.01. ANOVA test for FWHM of TIRF showed a significant difference (P < 0.01) only for *Nt*PIN11T-GFP. (*E*) Heterogeneity in the nanodomain distribution for three GFP-tagged *Nt*PINs, coefficient of average intensity variation from 40/30/47 induced cells of *Nt*PIN11T-GFP, *Nt*PIN2T-GFP and *Nt*PIN3bT-GFP. Letters indicate significantly different groups, one-way ANOVA with post-hoc Tukey’s honest significant difference test; P < 0.01. Error bars = SD. (*F*) PM ghost TIRFM images of anti-GFP immunostained tagged *Nt*PINs nanodomains. (*G*) Comparison of nanodomain dimensions imaged by TIRFM *in vivo* and in immunostained PM ghosts (FWHMs). Data from 20/12 (*In vivo*/PM ghosts) individual cells for each protein, n=582/889 (*In vivo*/PM ghosts, *Nt*PIN11T), 580/812 (*In vivo*/PM ghosts, *Nt*PIN2T), and 650/847 (*In vivo*/PM ghosts, *Nt*PIN3bT). (*H*) Comparison of coefficient of average intensity reflecting nanodomain distribution heterogeneity imaged by TIRFM in whole cells *in vivo* and immunostained PM ghosts. Data from 40/12 cells (*In vivo*/PM ghosts, *Nt*PIN11T), 30/8 (*In vivo*/PM ghosts, *Nt*PIN2T), and 47/13 cells (*In vivo*/PM ghosts, *Nt*PIN3bT). The significance of the differences in FWHM for TIRFM/SRRF (D) and *in vivo*/PM ghosts (*G*), and for coefficient of average intensity for *in vivo*/PM ghosts (*H*) were all tested with T-test; horizontal lines show the mean values, ***P < 0.01. Scale bars 2 (B), 3 (C), and 5 (F) µm.

To test if the PM localizations of individual *Nt*PINs are related to the cell wall context, we performed *in vivo* Airyscan confocal and TIRF microscopy (TIRFM) in the exponential cells and compared their nanodomain patterns with those from fixed PMs released from isolated protoplasts. In living cells, both *Nt*PIN2T-GFP and *Nt*PIN3bT-GFP showed clear signal variability along the z-axis and their nanodomains were spatially more discrete than for *Nt*PIN11T-GFP, where we observed a highly packed and fuzzy signal (Fig. 2*B*). We further supported these results by *in vivo* TIRFM imaging combined with the superresolution radial fluctuation (SRRF) algorithm (Gustafsson et al., 2016), which revealed similar patterns of nanodomains with full-width half-maximum (FWHM) around 200 nm for all *Nt*PINs (Figs. 2*C, D*). This is in agreement with dimensions of nanodomains reported previously from TIRFM-based imaging of PIN auxin carriers in Arabidopsis root and hypocotyl cells (McKenna et al., 2019; Narasimhan et al., 2021). Analysis of the variability of TIRFM fluorescence again showed that *Nt*PIN11T-GFP had the most homogeneous distribution within the PM (Fig. 2*E*). After the removal of the cell wall, nanodomains of all *Nt*PINs were still present (Fig. 2*F*), but they were larger for *Nt*PIN2T-GFP and *Nt*PIN3bT-GFP or unchanged in size for *Nt*PIN11T-GFP (Fig. 2*G*). However, the overall degree of homogeneity in the distribution decreased after the removal of the cell wall for all *Nt*PINs, including *Nt*PIN11T-GFP (Fig. 2*H*).

In conclusion, we have shown here that the removal of the cell wall resulted in a much lower homogeneity of signals for all NtPINs, suggesting that the cell wall defines their localization patterns within the PM.

### Electron microscopy reveals a homogenous distribution of *Nt*PIN11T in the PM in contrast to the other two *Nt*PINs

The level of spatial resolution of TIRFM cannot accurately reveal the nature of the different heterogeneity in the distribution of individual GFP-tagged *Nt*PINs. Therefore, we used two independent methods of electron microscopy to investigate the PM localization of *Nt*PINs in fixed PMs released from isolated protoplasts and in whole cells.

For PMs from isolated protoplasts, we used correlative light and electron microscopy (CLEM) (Stelate et al., 2021), using a combination of TIRFM of immunostained GFP-tagged *Nt*PINs and advanced low-voltage environmental scanning microscopy (A-ESEM). The positions of fluorescent nanodomains observed by TIRFM were spatially correlated with EM images after their mutual superimposition (Fig. 3A), which allowed us to quantify the number of gold particles corresponding to a fluorescent nanodomain for all GFP-tagged *Nt*PINs. While nanodomains with *Nt*PIN2-GFP and *Nt*PIN3b-GFP contained on average five to six individual PIN molecules, nanodomains with *Nt*PIN11-GFP contained significantly more individual molecules, on average three times as many (Fig. 3*B*). This result nicely supports the higher aggregation of the originally more diffuse fluorescence signal observed by TIRFM in *Nt*PIN11-GFP after the removal of the cell wall (Fig. 2*H*).

**Fig. 3.**
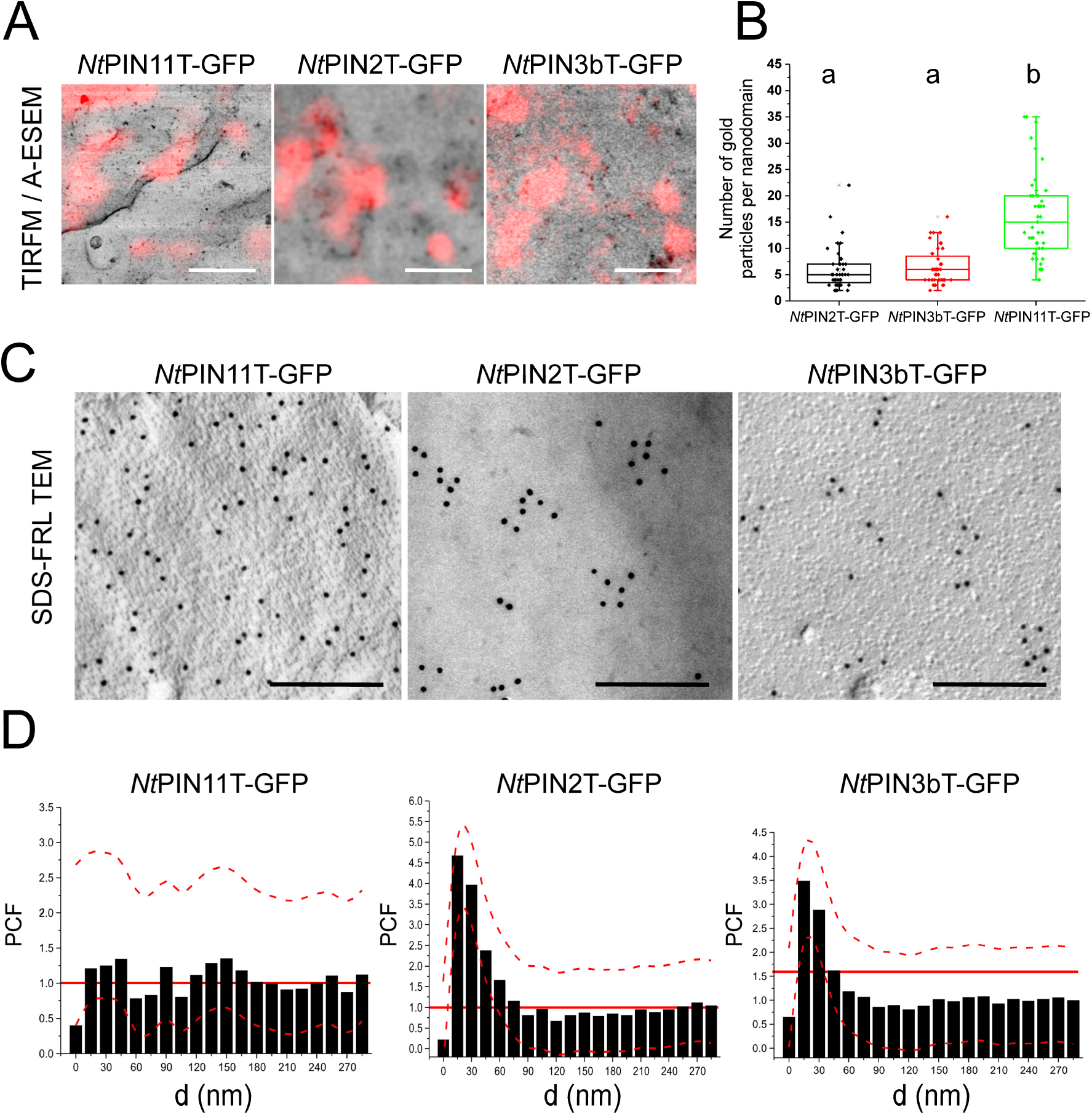
Distribution of individual *Nt*PINs revealed by EM. (*A*) TIRFM/A-ESEM CLEM images of anti-GFP immunostained PM ghosts generated from three GFP-tagged *Nt*PINs. Samples immunostained with anti-GFP primary polyclonal rabbit antibody and goat anti-rabbit IgG antibody conjugated with Alexa fluor 546 (red signals) and colloidal gold (1 nm) with 2-min gold enhancement (black dots). (*B*) The number of gold particles per one nanodomain spatially correlated with TIRFM/A-ESEM, the number of analyzed domains was 45 (*Nt*PIN11T-GFP), 44 (*Nt*PIN2T-GFP), and 45 (*Nt*PIN3bT). Letters indicate significantly different groups, one-way ANOVA with post-hoc Tukey’s honest significant difference test; P < 0.01. Error bars = SD. (*C*) High-pressure freezing SDS-FRL of *Nt*PINs in the P side of the tobacco cell PM. Platinum SDS-digested replicas labeled with polyclonal rabbit anti-GFP antibody followed by a goat anti-rabbit IgG antibody conjugated with colloidal gold (12 nm). (*D*) Quantification of gold interpoint distances (d) distribution in SDS-FRL images expressed as calculated pair correlation function (PCF) fitted to the equation described in materials and methods. The number of analyzed images (5×5 µm) was 14 (*Nt*PIN11T-GFP), 13 (*Nt*PIN2T-GFP), and 16 (*Nt*PIN3bT). Note the random distribution of *Nt*PIN11T-GFP, compared to the other two *Nt*PINs. Dashed red lines represent SD. Bold red lines represent the simulated random distribution of PCF. Scale bars 1 µm (*A*) and 250 nm (*C*).

Whole cells were subjected to high-pressure freezing and sodium dodecyl sulphate-digested freeze-fracture replica immunolabeling (SDS-FRL) protocol according to Fujimoto (1995). Quantitative image analysis revealed that *Nt*PIN11T-GFP was highly randomly distributed in the PM, in contrast to the aggregated distribution of *Nt*PIN2T-GFP and *Nt*PIN3bT-GFP (Figs. 3*C*, *D*). These results provided independent evidence for the differential localization of individual *Nt*PINs at the single molecule level. Importantly, the distribution of *Nt*PIN2T-GFP and *Nt*PIN3bT-GFP was very similar to the pattern observed for immunostained phosphatidylinositol 4,5-bisphosphate (PI(4,5)P2), which we used as a positive control for PM (*SI Appendix*, Fig. S3). The SDS-FRL method has been used in PM-related research in plants for studies of cellulose synthase complexes (Kimura et al., 1999) and more recently for the localization of Arabidopsis PIN2 (Li et al., 2021), yielding very similar patterns to those we show here for *Nt*PIN2T-GFP.

Overall, the use of CLEM and SDS-FRL revealed the existence of specific PM distribution patterns for individual PINs at the single molecule level and supported the more homogeneous nature of the *Nt*PIN11T distribution.

### Differential dynamics of *Nt*PINs within the PM

The different distributions of *Nt*PINs suggest that they may also be differentially restricted in their lateral PM mobility and compartmentalization, which has previously been shown in yeast to define the function and turnover of the methionine transporter (Busto et al., 2018). Our previous results showed that *Nt*PIN2T-GFP, *Nt*PIN3bT-GFP and *Nt*PIN11T-GFP are all localized in the transverse and lateral PMs in BY-2 cells (Müller et al., 2019). We detected them also in the developing cell plates from the very early stages of their formation (Fig. 4*A*). These localizations allowed us to perform a series of fluorescence after photobleaching (FRAP) experiments to test the dynamics of GFP-tagged *Nt*PINs in cell plates and mature transverse and lateral PMs (Fig. 4*B*). The results showed again a different behavior of *Nt*PIN11T-GFP, which was immobilized already in the growing cell plates (Fig. 4*B*, left panel), whereas the other two *Nt*PINs were significantly more dynamic. In mature transverse and lateral PMs, the immobile fractions of all *Nt*PINs were consistent with published data for integral PM proteins, including some PINs (Martinière et al., 2012; Martinière & Runions, 2013). The lower mobility of *Nt*PIN11T-GFP in the cell plate may reflect its specific retention already in the center of the growing cell plate, from where the other two PIN proteins are actively retrieved by the endocytosis (Mravec et al., 2011). To discriminate between the mobilities of nanodomains with individual NtPINs in the mature PMs, we used the TIRFM single particle tracking (SPT) to analyze the mean square displacement (MSD) and diffusion rates of nanodomains. This allowed us to show that even in mature lateral PMs, nanodomains with *Nt*PIN11T-GFP were the most restricted in their mobility, in contrast to nanodomains with *Nt*PIN2-GFP, which were the most mobile (Figs. 4*C*-*E*).

**Fig. 4.**
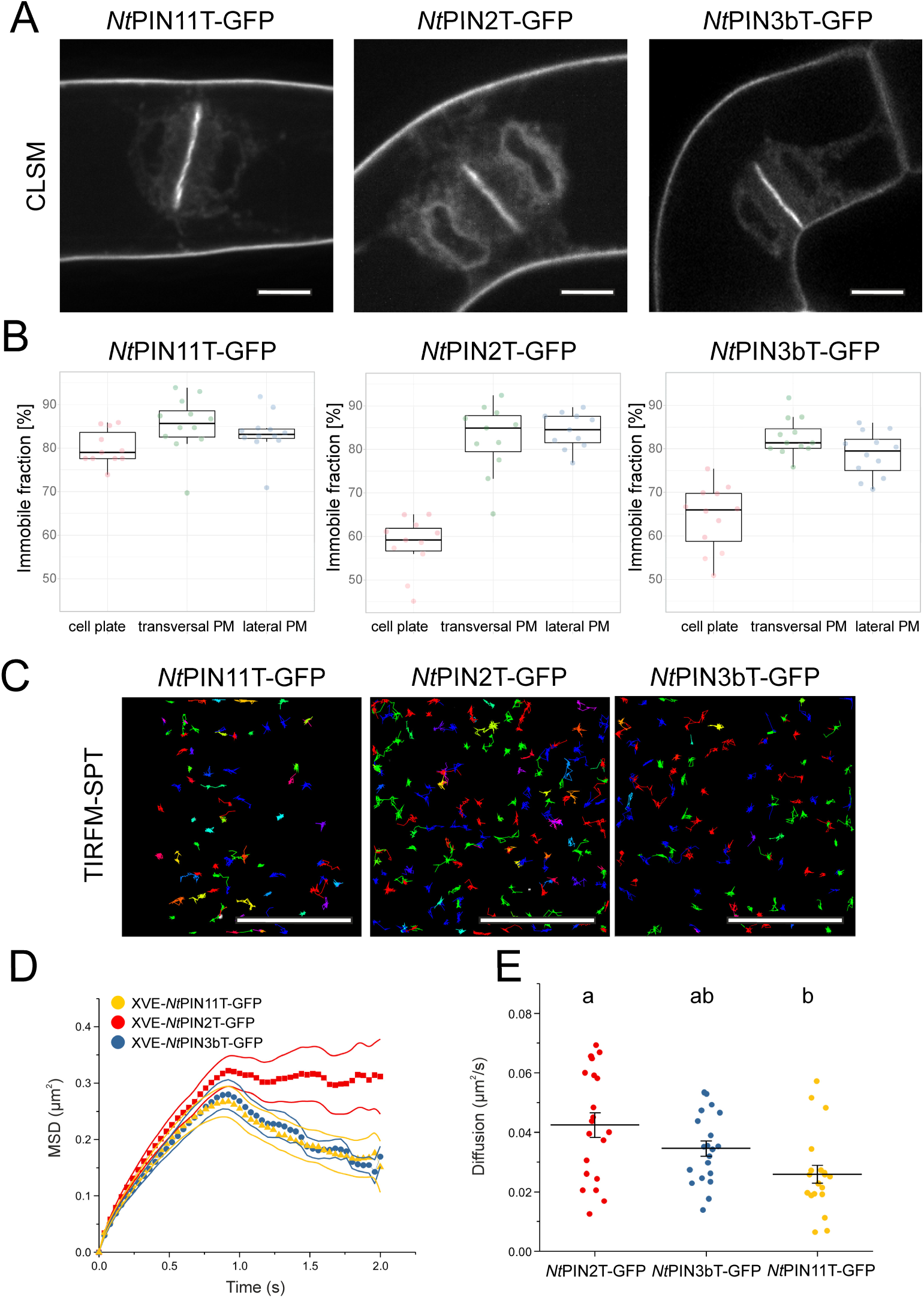
Differential dynamics of *Nt*PINs within PM. (*A*) CLSM of 2-day-old induced *Nt*PIN11T-GFP, *Nt*PIN2T-GFP and *Nt*PIN3bT-GFP cells showing the signal at the cell plates and mature PM. Single confocal sections. (*B*) FRAP analysis of three GFP-tagged *Nt*PINs in cell plates and transverse and lateral PMs. Note remarkably higher dynamics of *Nt*PIN11T-GFP in cell plates, represented by a higher percentage of the immobile fraction of molecules that are not recovering during FRAP recovery period. (*C*) Representative images of trajectories of individual GFP-tagged *Nt*PIN molecules acquired with TIRFM in 2-day-old induced cells, detected with SPT algorithm. Trajectories represent 2 types of dynamics, i.e. constrained movements (shown with short trajectories) and free diffusion (shown with long trajectories). (*D*) MSD calculated from TIRFM images. The number of evaluated trajectories was 989 (*Nt*PIN11T-GFP), 281 (*Nt*PIN2T-GFP), and 847 (*Nt*PIN3bT-GFP). Note more constrained dynamics of *Nt*PIN11T-GFP and *Nt*PIN3T-GFP compared to *Nt*PIN2T-GFP. The thick bold lines represent the standard error. (*E*) Diffusion rates of GFP-tagged *Nt*PINs in the PM (from left to right trajectories from 23, 20, and 23 cells analyzed). Note the lower diffusion of *Nt*PIN11T-GFP in contrast to *Nt*PIN2-GFP. Letters indicate statistically significant differences between groups, one-way ANOVA with post-hoc Tukey’s honest significant difference test, P < 0.01). Error bars indicate standard error. Scale bars 10 µm.

In summary, the study of the dynamics of *Nt*PINs in the PM revealed their distinct behavior and highlighted the remarkable stability of *Nt*PIN11T-GFP already in the developing cell plates.

### *Nt*PINs interact similarly with anionic phospholipids in molecular dynamics simulations

To test if differential direct interactions with anionic phospholipids cause the observed differences in distribution and mobility between *Nt*PINs, we first examined the structures of *Nt*PIN homodimers predicted by Alphafold2 with very high confidence (Fig. 5*A*; *SI Appendix*, Fig. S4*A*) in agreement with the recently resolved experimental structures of several different Arabidopsis PIN proteins organisms (Su et al., 2022; Ung et al., 2022; Yang et al., 2022). Mapping Coulombic electrostatic potential on the structures of distinct *Nt*PIN dimers did not show any pronounced differences in the position of positively charged amino acids, which could mediate direct contact with anionic phospholipids of the plasma membrane (Fig. 5*B*). Next, we used the coarse-grained molecular dynamics (CG-MD) simulation, a widely used computational method to investigate protein-lipid interactions that can reveal so-called lipid fingerprints characteristic of any integral membrane protein (Corradi et al., 2018; Neubergerová and Pleskot, 2024). To this end, we embedded the *Nt*PIN dimers into a proxy for the asymmetrically charged plant PM. CG-MD simulations were performed without long disorder cytosolic loops predicted with very low confidence (Fig. 5*A*; *SI Appendix*, Fig. S4*A*) as these unstructured parts would hamper modeling convergence. Five CG-MD replicas performed in total for 25 μs for each *Nt*PIN pair did not reveal any substantial differences in the lipid fingerprints. Additionally, we did not observe any differences in the protein diffusion through the lipid bilayer (Fig. 5*C*).

**Fig. 5.**
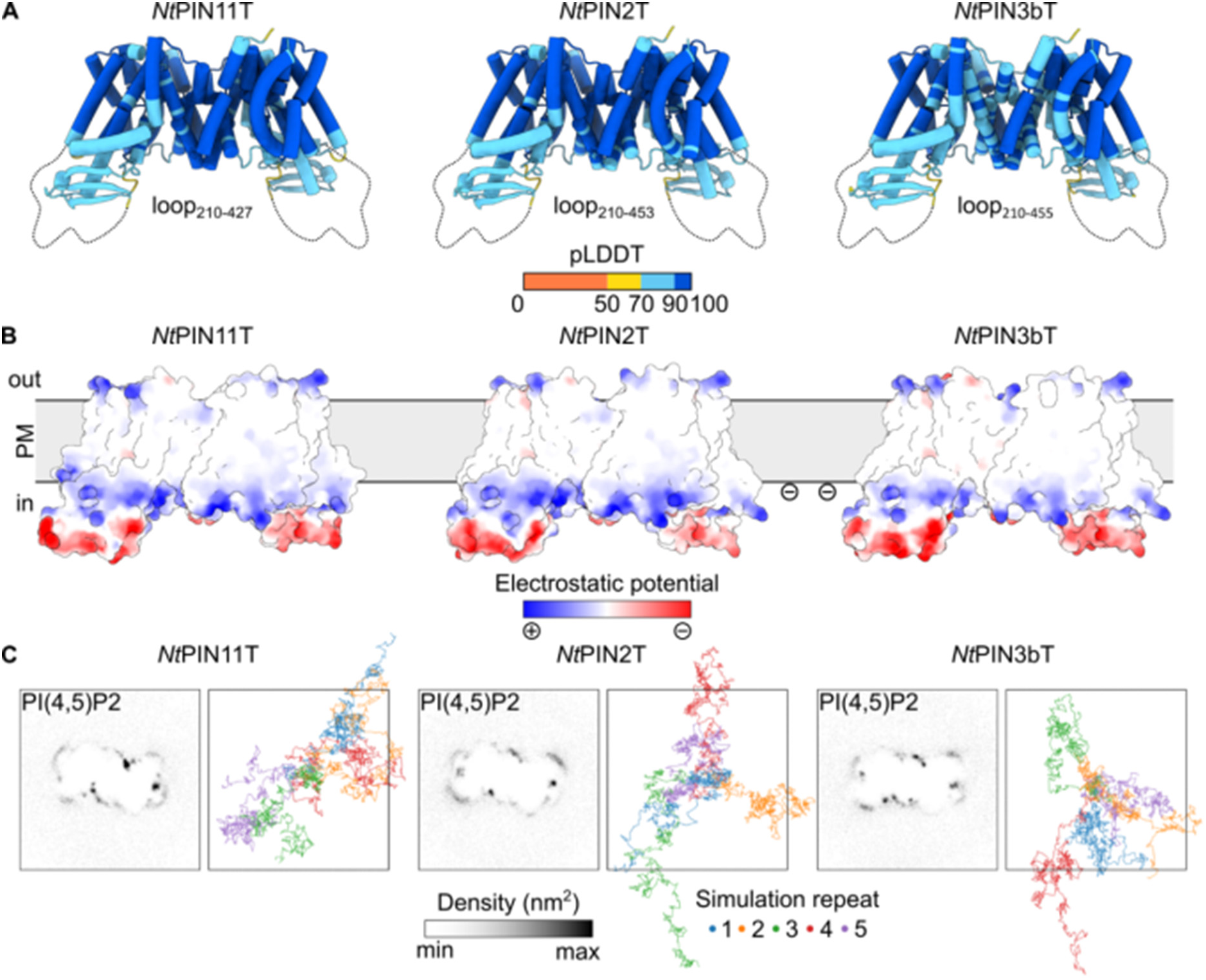
Protein-lipid interactions of *Nt*PINs. (*A*) AlphaFold2 predictions of transmembrane regions of *Nt*PIN11T, NtPIN2T and *Nt*PIN3bT dimers. The loop, predicted with the low pLDDT score, is visualized only schematically. (*B*) Electrostatic potential mapped on transmembrane regions of *Nt*PIN11T, *Nt*PIN2T and *Nt*PIN3bT dimers. All *Nt*PINs have positively charged patches located on the cytoplasmic side of the membrane possibly interacting with negatively charged anionic phospholipids. PM = plasma membrane. (*C*) PI(4,5)P2 densities around transmembrane regions of *Nt*PIN11T, *Nt*PIN2T and *Nt*PIN3bT dimers (left) and protein mobility in the model plasma membrane (right) based on molecular dynamics simulations. No significant difference in PI(4,5)P2 clustering pattern nor protein mobility were observed among *Nt*PINs dimers.

Altogether, our computational data suggest that the differences between distinct *Nt*PIN isoforms are not caused by differential protein-lipid interactions.

### Specific interactions of *Nt*PIN11 with cell wall suggested by pharmacology treatments

The dynamics, localization, and function of PIN proteins are determined not only by their interaction with specific PM phospholipids (Li et al., 2020; Marhava, 2022) but also by their interaction with the cell wall and cytoskeleton (Ke et al., 2021; Kleine-Vehn et al., 2008; Martiniere et al., 2012). Therefore, we used a pharmacological approach combined with *in vivo* experiments to understand which cellular structures might stay behind the differential mobility and distribution of three *Nt*PIN proteins. We have analyzed MSD and diffusion rates, which revealed significantly increased mobility of *Nt*PIN11T-GFP after the inhibitor of cellulose biosynthesis isoxaben (Tateno et al., 2016), as well as by the application of actin drug latrunculin B or microtubule drug oryzalin and their combinations (Fig. 6*A*). Drug-induced increased mobility of *Nt*PIN11T-GFP may reflect the loss of natural interactions within the crosslinked network of the cortical cytoskeleton and cell wall (Feraru et al., 2011; McKenna et al., 2019; Tolmie et al., 2017). Interestingly, both cytoskeleton and cell wall inhibitors did not change the mobility of *Nt*PIN2T-GFP (Fig. 6*B*) and there was only a weak effect of oryzalin on the mobility of *Nt*PIN3bT-GFP (Fig. 6*C*). In addition, we did not observe any significant changes in the dimensions of nanodomains (Fig. 2G) after drug treatments for *Nt*PIN11T-GFP in contrast to their increased size in *Nt*PIN3bT-GFP (Fig. 6C) and *Nt*PIN2T-GFP (Fig. 6B) cells. These results independently confirm the unchanged dimensions of *Nt*PIN11T-GFP nanodomains in PM ghosts (Fig. 2*G*). Finally, we investigated whether the inhibitors interfered with auxin efflux in induced *Nt*PIN11T-GFP, *Nt*PIN2T-GFP and *Nt*PIN3bT-GFP cells. For cytoskeletal drugs, we observed a decrease in the accumulation of [^3^H]NAA in all three GFP-tagged *Nt*PIN lines (Fig. 6*D*) to the same levels as in drug-untreated controls (Fig. 2*A*). However, there was a significant auxin efflux-promoting activity of isoxaben in induced *Nt*PIN11T-GFP cells, in contrast to *Nt*PIN2T-GFP and *Nt*PIN3bT-GFP (Fig. 6*E*).

**Fig. 6.**
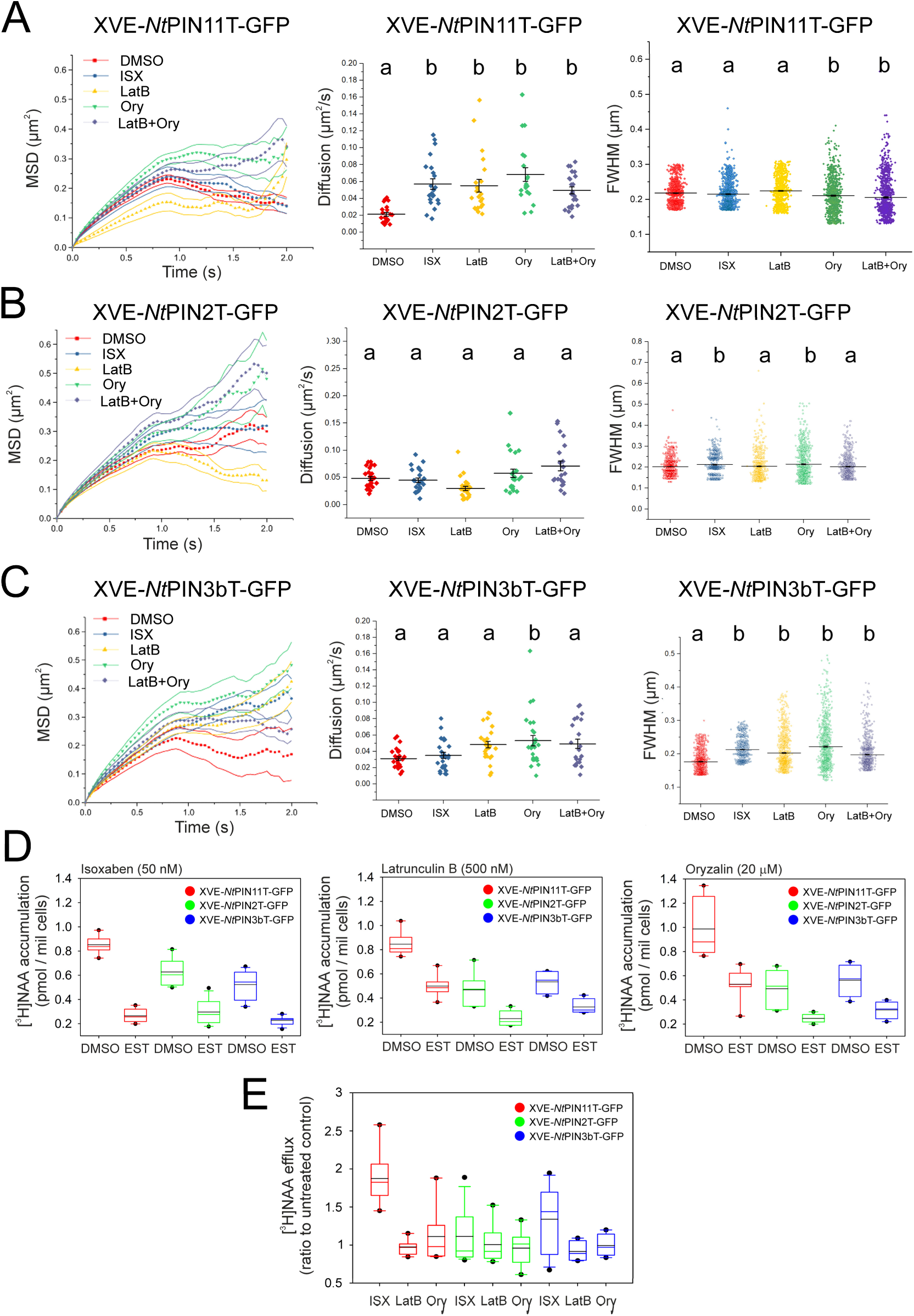
Cytoskeleton and cell wall drugs affect the dynamics, nanodomain organization, and function of *Nt*PINs. (*A*-*C*) Mean square displacement (MSD) curves (left columns), diffusion rates (middle columns), and individual nanodomain dimensions (right columns) for *Nt*PIN11T-GFP (*A*), *Nt*PIN2T-GFP (*B*) and *Nt*PIN3bT-GFP (*C*), calculated from TIRFM images. Treatments with mock (DMSO), 10 µM isoxaben, 500 nM latrunculin B, or 20 µM oryzalin and their combinations. MSD for *Nt*PIN11T-GFP, n=940 (DMSO), 1006 (ISX), 991 (LatB), 892 (Ory), 1025 (LatB+Ory), *Nt*PIN2T-GFP, n=1017 (DMSO), 1020 (ISX), 989 (LatB), 950 (Ory), 889 (LatB+Ory), and *Nt*PIN3bT-GFP, n=1045 (DMSO), 1007 (ISX), 1012 (LatB), 1020 (Ory), 985 (LatB+Ory). Diffusion rates for *Nt*PIN11T-GFP, calculated from left to right from 21, 22, 22, 20, and 23 cells, for *Nt*PIN2T-GFP, calculated from left to right from 25, 25, 25, 23, and 21 cells, and for *Nt*PIN3bT-GFP, calculated from left to right from 27, 25, 26, 26, and 22 cells. Nanodomain FWHM dimensions, for *Nt*PIN11T-GFP, calculated from left to right from 884, 893, 894, 815, and 898 nanodomains, *Nt*PIN2T-GFP, calculated from left to right from 935, 896, 803, 876, and 891 nanodomains, and *Nt*PIN3bT-GFP, calculated from left to right from 1059, 1062, 1057, 1002, and 1045 nanodomains. Letters indicate statistically significant differences between groups, one-way ANOVA with post-hoc Tukey’s honest significant difference test; P < 0.01. Error bars = standard error. (*D*) Analysis of effects of cytoskeletal and cell wall drugs on auxin transport in XVE-*Nt*PIN11T-GFP, XVE-*Nt*PIN2T-GFP, and XVE-*Nt*PIN3bT-GFP lines induced with 1 µM β-estradiol for 48 h (EST). 50 nM isoxaben (left), 500 nM latrunculin B (middle), and 20 µM oryzalin (right). [^3^H]NAA accumulations from six biological repetitions (independent inductions in the particular line) are shown with a mean (black lines) and median (color lines). Data points outside 1.5 times interquartile range from 1st/3rd quartile are shown as outliers. The significance at P < 0.05 was tested positively with the Mann-Whitney U-test for all induced versus non-induced cells. (*E*) [^3^H]NAA efflux expressed as a ratio of mean values of [^3^H]NAA accumulations in non-induced and induced cells after drug treatments expressed as ratios of drug-treated cells to drug-untreated controls.

These results suggest that the interaction with the cell wall is important for the stabilization of the *Nt*PIN11 in the PM and this stabilization modulates its auxin efflux function.

### Identifying specific interactors of *Nt*PINs using co-IP/MS^2^

Considering variable localization and dynamics of each *Nt*PIN, it can be postulated that these properties are a consequence of their specific interactions (Marhava, 2022). Unfortunately, the latest research does not yet yield sufficient results from biochemical analyses specifically aimed at identifying PIN-interacting proteins in the PM. Available are anti-GFP co-IP data from whole Arabidopsis plants carrying pPIN1::PIN1-GFP, which revealed an important role for the dynamin DRP1A in PIN polarization and endocytosis (Mravec et al., 2011), and some as yet untested PIN interactor candidates have been suggested by yeast membrane-bound interactome (Jones et al., 2014). In contrast to previously published more general co-iPs on PINs, we rather focused primarily on identifying membrane-related interactors. Our anti-GFP immunoblot analysis using solubilized membrane fractions from 2-day-old induced and non-induced cells showed the presence of a distinct band corresponding in size to individual *Nt*PIN-GFPs and several degradation products, while there was no signal detected in non-induced cells (*SI Appendix*, Fig. S5). Eluted co-immunoprecipitated fractions were trypsin-digested and analyzed by LC-MS^2^. Label-free relative quantification provided sets of peptides that we blasted against the tobacco genome database (Edwards et al., 2017), identifying the closest homologs using blast against the *Arabidopsis thaliana* proteome. In comparison with non-induced cells, we observed a significant enrichment of previously described PIN endocytosis-related interactors (*SI Appendix*, Tab. S4). To reveal specific and potentially novel interactors, we employed a rigorous filtering approach, excluding any potential interactors that were also identified, albeit to a minimal extent, in non-induced cells as well as in GFP-expressing cells (see Materials and Methods). Upon application of these strict filtering conditions, we obtained a novel interactor candidates for *Nt*PIN11T-GFP, *Nt*PIN2T-GFP, and *Nt*PIN3bT-GFP (Fig. 7A; *SI Appendix*, Tabs. S1-S3).

**Fig. 7.**
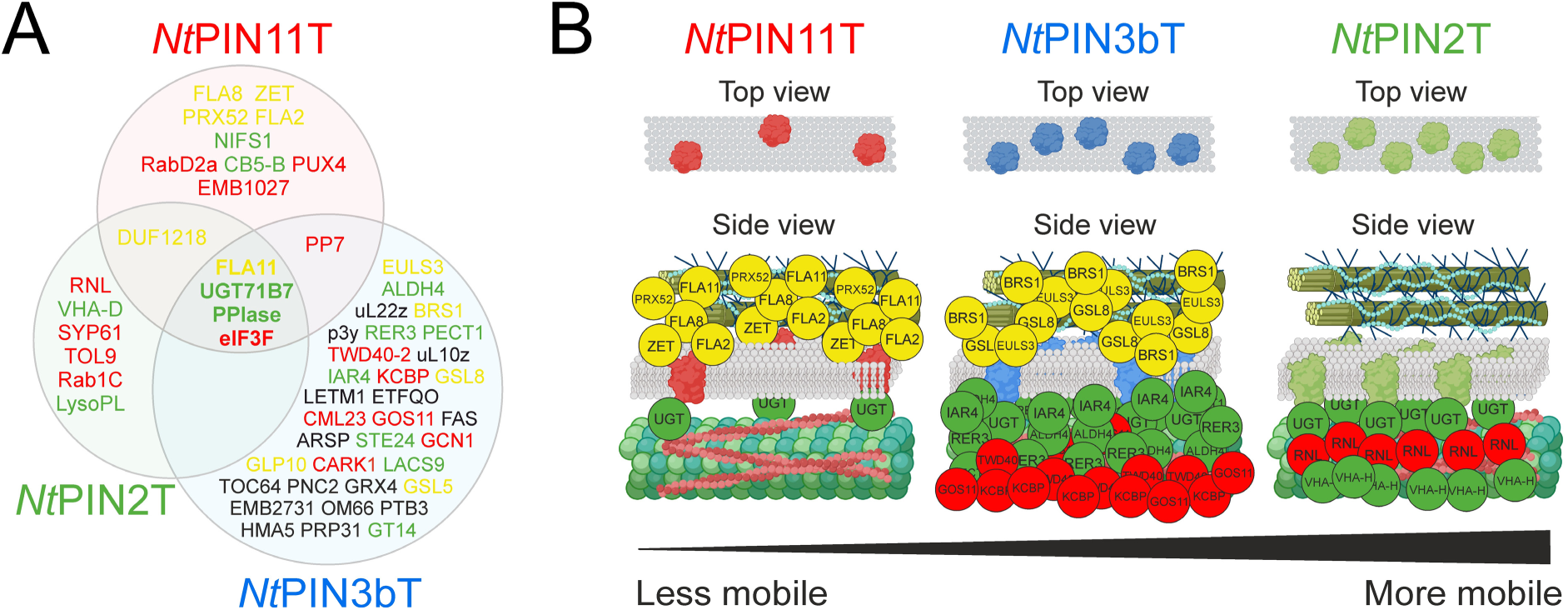
Candidate interactors of *Nt*PIN11T, *Nt*PIN2T, and *Nt*PIN3bT. (*A*) Venn diagram of the common and specific co-iP interactors for *Nt*PINs. Shown are all interactors whose abundance was increased at least 32-fold compared to the abundance of proteins with the lowest positively detected levels. Common interactors include fasciclin-like protein (FLA11), UDP-dependent glycosyl transferase (UGT71B7), cyclophilin-like peptidyl-prolyl cis-trans isomerase (PPIAse) and eukaryotic translation initiation factor 3 subunit F (eIF3). (*B*) Schematic representation of the results of localization and mobility tests of *Nt*PINs and the co-IP interactor screen. Shown are all interactors whose abundance was increased at least 128-fold compared to the abundance of proteins with the lowest detected levels. Note the preferential cell wall protein interactions for the more homogeneously distributed and less mobile *Nt*PIN11T in contrast to the preferential cytosolic interactions for the more aggregated and mobile *Nt*PIN2T. Interactors are color-coded as follows: intracellular signaling and vesicle trafficking proteins (red), cytosolic and membrane-bound enzymes (green), cell wall proteins (yellow), and other proteins including ribosomal proteins and membrane carriers (black). The abbreviations for every protein detected are described in *SI Appendix* Tabs. S1-S3. Graphical elements in B were created with BioRender.com.

Based on their cellular functions, we classified candidate interactors into three groups as shown in Fig. 7*A*, namely proteins involved in signaling and vesicle trafficking processes (red), cytosolic and membrane-bound enzymes (green), and cell wall proteins (yellow). In the group of signaling and vesicle trafficking processes, we identified a single common interactor, a homolog of the heterotrimeric GTPase eukaryotic translation initiation factor 3 subunit p47 (EIF2). EIF2 has been reported to co-fractionate with TPLATE (McWhite et al., 2020), part of the complex previously shown to function in the endocytosis of Arabidopsis PINs (Gadeyne et al., 2014). Moreover, also two of the *Nt*PIN-specific interactors (*Nt*PIN2T) were previously reported to be connected with auxin-regulated development. In particular, the tRNA ligase RLG1 has been implicated in tRNA maturation-associated auxin responses (Leitner et al., 2015), and the SNARE protein TOL9 has been shown to act in the trafficking of ubiquitinated PIN2 cargo (Korbei et al., 2013). In the group of cytosolic and membrane-bound enzymes, we identified two common interactors. First, homologs of cyclophilin-like peptidyl-prolyl cis-trans isomerase (PPIase), which was previously shown to catalyze cis/trans isomerization of conserved proline-containing peptides in *Arabidopsis thaliana* PIN1 changing their localization and activity possibly through specific phosphorylation (Xi et al., 2016). Cyclophilin A from tomato was also demonstrated to be involved in the PM localization of PINs (Ivanchenko et al., 2015). Second common interactor, tobacco homolog of *Arabidopsis thaliana* UGT71B7 UDP-glucosyl transferase *Nt*UGT3T (Müller et al., 2020) was previously documented to catalyze the glycosylation of oxIAA (Tanaka et al., 2014). Finally, in the group of cell wall proteins, we identified a single common interactor, a homolog of the *Arabidopsis thaliana* fasciclin-like arabinogalactan protein FLA11 (Ma et al., 2022) and two other homologs of FLA2 and FLA8, which were specifically found among the *Nt*PIN11T interactors, in addition to several other cell wall-related proteins (Fig. 7, *SI Appendix*, Tabs. S1-S3). FLA proteins are apoplastic glycosylphosphatidylinositol-anchored proteins that connect the PM and the cell wall and play role in the cellulose biosynthesis (Liu et al., 2020). They have been previously co-fractionated with PINs and ABCBs (Titapiwatanakun et al., 2009), which suggests that could therefore be part of the auxin-transporting domain within the PM-cell wall nexus (Feraru et al., 2011).

Altogether, the *Nt*PIN interactor screen provided an independent biochemical support for our microscopic observations, supporting the conclusion that the stronger association with cell wall may account for the *Nt*PIN11T homogenous distribution within PM.

## Conclusion

Our data reveal a previously unidentified phenomenon regarding the localization, dynamics, and function of auxin carriers, integral membrane proteins. To draw meaningful conclusions, it was essential to utilize a simplified plant cell model where the properties of individual proteins could be compared in an identical biological context. By employing tobacco cells expressing fluorescent fusions of three canonical tobacco auxin efflux carriers under inducible system, we demonstrated that within the group of auxin carriers, the homolog of *Arabidopsis thaliana* PIN1, *Nt*PIN11T, is the most homogenously distributed and the least mobile *Nt*PIN.

Its immobilization depends on the actin and microtubular cytoskeleton as well as the cell wall composition, which restricts its function. Moreover, our biochemical analysis supported the hypothesis that auxin transport proteins in the PM are in dynamic association not only with proteins ensuring their intracellular trafficking and interaction with the cell wall, but also with the enzymatic machinery of auxin metabolism. While PM aggregation of the particular PIN and its mobility correlates positively with the amount of cytosolic interactions, more homogenous distribution depends on cell wall protein interactions (Fig. 6*B*). This discovery needs to be supported by further studies using model plants.

## Materials and methods

### Plant material, culture conditions, drug treatments, and chemicals

Tobacco cell line BY-2, *Nicotiana tabacum* L. cv Bright Yellow 2 (Nagata et al., 1992) was cultured in liquid Murashige and Skoog (MS) medium (4.3 g/L MS salts, Sigma M5524) supplemented with 3% sucrose (w/v), 100 mg/L myo-inositol, 1 mg/L thiamine, 0.2 mg/L 2,4-D, and 200 mg/L KH_2_PO_4_, pH 5.8, in the darkness, at 27°C, under continuous shaking (150 rpm; orbital diameter 30 mm). Suspension-cultured cells were subcultured every 7 days. Stock calli cultured on the MS media solidified with 0.6% (w/v) agar were subcultured monthly. Transgenic cells and calli were maintained on the MS medium supplemented with 100 µg/mL cefotaxime and either 40 µg/mL hygromycin for XVE-*Nt*PIN11T-GFP, XVE-*Nt*PIN2T-GFP and XVE-*Nt*PIN3bT-GFP cells (Müller et al., 2019; Stelate et al., 2021) or 100 µg/mL cefotaxime for *Nt*PIN11T::*Nt*PIN11T:GFP, *Nt*PIN2T::*Nt*PIN2T:GFP, and *Nt*PIN3bT::*Nt*PIN3bT:GFP cells.

To induce an estrogen receptor-based chemical-inducible system (Zuo et al., 2000), MS medium was supplemented with 1 µM β-estradiol (10 mM stock solution in DMSO, Sigma-Aldrich Inc., St. Louis, MO, USA) at the beginning of the subculture interval. An appropriate amount of DMSO was added to the non-induced cells. For microscopy analyses and auxin transport assays, stock solutions of latrunculin B (2.53 mM), oryzalin (57.7 mM), and isoxaben (1 mM) in dimethyl sulfoxide (DMSO) were added to 4 ml aliquots of 2-day-old cell suspensions in a 6-well plate (microscopy) or 30 ml of 2-day-old cell suspensions in 100 ml flask (auxin transport assays), achieving final concentrations of 0.5 µM latrunculin B, 20 µM oryzalin, and 10 µM (microscopy) or 50 nM isoxaben (auxin transport assays). For all drug treatments in microscopy, cells were incubated for 30 min under constant shaking (80 rpm; orbital diameter 10 mm). For the auxin transport assay, cells were pre-treated with drugs for 2 h before the accumulation assays. An appropriate volume of DMSO was added for the mock treatment. Unless stated otherwise, all chemicals were supplied by Sigma Aldrich (St Louis, MO, USA).

### Cloning of tobacco PIN promoters, preparation of gene constructs and transgenic tobacco cells

For individual *NtPIN* genes, DNA promoter sequences for *NtPIN11T* (GenBank a.n. KC433528,-1500 bp from ATG) *NtPIN2T* (GenBank a.n. KP143727,-2069 bp from ATG) and *NtPIN3bT* (GenBank a.n. KC425459,-2000 bp from ATG) were amplified with PCR and cloned using GoldenBraid 2.0 cloning system (Sarrion-Perdigones et al., 2013), as previously described (Müller et al., 2021). They were used to replace inducible promoters in GFP-tagged *NtPIN* genes (Müller et al., 2019). Transformation of BY-2 cells was done by co-cultivation with *Agrobacterium tumefaciens* strain GV2260 as described previously (Petrášek et al., 2003). Transgenic lines were harvested after 4 weeks, cultured on solid media with kanamycin, and individual suspensions established in multiwell plates were observed using fluorescence wide-field and confocal microscopy. At least 40 individual lines were screened for every construct.

### RNA-seq analysis and RT-qPCR

RNA-seq analysis for the comparison of dividing and elongated cells was performed according to a previously published protocol (Müller et al., 2021). Briefly, total RNA was isolated from 50 to 100 mg of cells using RNeasy Plant Mini kit (Qiagen, Hilden, Germany) and treated with DNA-Free kit (Thermo Fischer Scientific, Waltham, MA, USA). RNA purity, concentration, and integrity were evaluated on 0.8% agarose gels (v/w) and by the RNA Nano 6000 Assay Kit using the Bioanalyzer (Agilent Technologies, Santa Clara, CA, USA). Approximately 5 mg of RNA isolated from 7 samples from 2-day-old and 6 samples from 7-day-old cells were analyzed by Eurofins (Ebersberg, Germany), providing at least 20 million 50 bps long reads. Data were deposited in NCBI (GSE160438, GSE275968).

RT-qPCR analysis was performed as described in detail in our previous works (Müller et al., 2019, 2020), including the method for the determination of relative transcript levels. 1 mg of DNAse-treated RNA isolated from samples harvested during life or cell cycle, were reverse-transcribed using M-MLV reverse transcriptase, RNase H-, point mutant (Promega Corporation, Madison, WI, USA). Quantitative real-time PCR was performed using GoTaq qPCR Master Mix (Promega Corporation, Madison, WI, USA) at 58°C annealing temperature on a LightCycler480 instrument (Roche, Basel, Switzerland). Tobacco EF1a (GenBank AF120093) was used as a reference for the relative quantification of the expression. At least three biological replicates with at least two technical repetitions for each were performed.

### Cell cycle synchronization

Synchronization of tobacco BY-2 was performed as described previously (Kuthanova et al., 2008). 15 µM aphidicolin, a reversible inhibitor of DNA polymerase α, was added to freshly inoculated stationary 7-day-old cells for 24 hours. After washing cells with fresh medium, mitotic index was scored during the progression of cell cycle after staining cells with Hoechst 33258 (Thermo Fischer Scientific, Waltham, MA, USA). At least 500 cells were counted in each of sampling time using a fluorescence microscope.

### Auxin transport assays

Determination of auxin efflux capacity of induced and non-induced XVE-*Nt*PIN11T-GFP, XVE-*Nt*PIN2T-GFP and XVE-*Nt*PIN3bT-GFP cells was performed with previously described protocol utilizing highly lipophilic [^3^H]NAA (naphthalene-1-acetic acid; 20 Ci/mmol; American Radiolabelled Chemicals, Inc., St. Louis, MO, USA) as the tracer for auxin efflux (Delbarre et al., 1996; Petrášek & Zažímalová, 2006). Briefly, 1 µM β-estradiol-induced or non-induced cells were grown for 2 days, filtered using a 20-µm nylon mesh, resuspended in uptake buffer (20 mM MES, 40 mM sucrose, 0.5 mM CaSO_4_; pH 5.7) and equilibrated for 45 min under continuous shaking. After washing in fresh uptake buffer, cell densities were adjusted to around 7. 10^5^ cells/mL after counting the cells in the Fuchs-Rosenthal hemocytometer, and cells were incubated for an additional 90 min in the darkness at 25°C. Accumulation assays were performed with [^3^H]NAA added to a final concentration of 2 nM. 0.5 mL aliquots were collected during 10 min accumulation period by filtration under reduced pressure using cellulose filters. Filters with cell cakes were transferred to scintillation vials and extracted for 30 min with 96% (v/v) ethanol. After the addition of a scintillation cocktail for liquid samples (EcoLiteTM, MP Biomedicals, Santa Ana, CA, USA) and 30 min rigorous shaking, radioactivity was determined by liquid scintillation counting with automatic correction for quenching (Packard Tri-Carb 2900TR scintillation counter; Packard Instrument, Fallbrook, CA, USA). DPM data were converted to pmols of [^3^H]NAA per 1 million cells and values were expressed at the time 10 min after the addition of [^3^H]NAA, separately for β-estradiol-induced cells and DMSO-treated non-induced controls. The resulting values are from at least six biological repetitions. Statistic evaluation (Sigma Plot, Systat software, Inc.) testing the significance with Mann-Whitney U-test for induced versus non-induced cells is provided in the particular plot.

### Preparation and immunostainings of PM ghosts

The preparation of PM ghosts was performed as described previously in detail (Stelate et al., 2021). Briefly, protoplasts were isolated from 2-day-old cells using a solution of 1% cellulase “Onozuka” R-10 (Yakuruto Honsha Co., Ltd., Tokyo, Japan) and 0.1% pectolyase Y-23 (Kyowa Chemical Products Co., Ltd., Osaka, Japan) in 0.45 M mannitol, following the previously published protocol (Krtková et al., 2012; Sonobe & Takahashi, 1994). They were collected and plated for 3 min on coverslips (10 mm Ø, thickness n. 1.5H, 170 µm± 5 µm; Paul Marienfeld GmbH & Co. Lauda-Königshofen, Germany) coated with poly-L-lysine. Membrane ghosts were generated by several quick flicks of the coverslip. PM ghosts adhering to the coated coverslips were fixed for 1 hour in a solution containing 4% paraformaldehyde, which was prepared from an aqueous 32% paraformaldehyde solution of EM grade (Electron Microscopy Sciences, Hatfield, PA, USA).

Immunostaining of protoplast ghosts was performed in a moisture chamber with parafilm on the bottom, as specified previously in detail (Stelate et al., 2021). The coverslips were incubated overnight at 4°C in one drop (20 µL) of a primary rabbit polyclonal anti-GFP antibody (cat. no. AS152987, Agrisera AB, Vännäs, Sweden) diluted 1:1000 in blocking solution with 0.5% bovine serum albumin (BSA) in phosphate-buffered saline (PBS). After washing three times in PBS, the coverslips were incubated for 45 min at room temperature with Alexa Fluor® 546-FluoroNanogold IgG goat anti-rabbit IgG secondary antibody (Cat. No. 7403, Nanoprobes, Yaphank, NY, USA) diluted 1:200 in 0.5% BSA. To increase the size of the gold particles, the coverslips were subsequently incubated for 2 min in a gold enhancement reaction solution (GoldEnhanceTM EM plus mixture, cat. no. 2114, Nanoprobes, Yaphank, NY, USA), prepared according to (Stelate et al., 2021). These preparations were stored in a moisture chamber until microscopy was performed.

### Fluorescence microscopy and laser scanning confocal microscopy

For routine *in vivo* microscopy inspection of induced cells and screening of fluorescence during the selection of new transgenic lines, upright Nikon Eclipse E600 microscope (Nikon, Tokyo, Japan) was used.

*In vivo* confocal laser scanning microscopy was performed with Zeiss LSM 880 inverted confocal laser scanning microscope (Carl Zeiss, Jena, Germany) using water immersion C-Apochromat objective 40x (NA 1.2). Fluorescence was excited with Argon 488 nm laser and GFP emission between 490 and 550 nm was recorded using GaAsP-PMT detectors or standard PMT detectors. Single confocal sections or maximum intensity projections of z-stacks were created using ZEN black software (Carl Zeiss, Jena, Germany). For Airyscan imaging, oil immersion Alpha Plan-Apochromat objective 100x (M27, NA 1.46) was used in combination with the array of GaAsP-PMT area detectors achieving the improvement of the spatial resolution to 120/120/350 (X/Y/Z nm). FRAP analysis was performed using FRAP module implemented in Zeiss Zen software. Rectangular regions (5×2 µm) were bleached within 6-10 s and images were acquired during the post-bleach period with a high frame rate to cover the time period 90-120 s. Values of immobile fraction were evaluated using a single exponential fit, calculated using FRAP module implemented in ZEISS Zen software, according to the previously described method (Laňková et al., 2016).

### TIRFM imaging

TIRFM of living cells and immunostained PM ghosts was performed in cells induced with 1 µM β-estradiol during the inoculation and cultured for two days. Zeiss ELYRA PS.1 workstation (Carl Zeiss, Jena, Germany), mounted on a fully motorized Axio Observer inverted microscope, was used with oil immersion Alpha Plan-Apochromat objective 100x (DIC M27 Elyra, NA 1.46, WD 0.11 mm), Immersol 518 F for 30°C immersion oil, and tube lens 1.6x.

Living cells were imaged in MS medium on a coverslip (10 mm Ø, thickness n. 1.5H, 170 µm± 5 µm; Paul Marienfeld GmbH & Co. Lauda-Königshofen, Germany). GFP was excited with 488 nm diode laser (300 mW; tuned to 0.6%), and fluorescence was recorded with the PCO Edge 5.5 scientific complementary metal-oxide-semiconductor (sCMOS) camera using the BP 495-550+LP750 beam splitter (16-bit, pixel size 40 × 40 nm).

Fixed PM ghosts adhered to the coverslip (10 mm Ø, thickness n. 1.5H, 170 µm± 5 µm; Paul Marienfeld GmbH & Co. Lauda-Königshofen, Germany) were covered with a small drop of dH_2_O, and a bright-field image of the entire slide was acquired with the Plan-Apochromat 10x objective (NA 0.45, WD 2 mm). The PM ghosts with the flattest appearance were selected, and their XY coordinates were saved for subsequent TIRFM as well as subsequent A-ESEM. Imaging individual PM ghosts was performed at selected positions with the 100x objective described above. Alexa Fluor 546 was excited with a diode-pumped solid-state laser at 561 nm (200 mW; tuned to 0.4 %), and the signal was collected with the sCMOS camera using the BM570-620+LP750 beam splitter (16-bit, pixel size 60 × 60 nm). After the acquisition, the coverslips were carefully removed, placed individually in the multiwell plate, and dried for 48 hours at RT for A-ESEM. For further details on the specimen holder for TIRFM/A-ESEM CLEM please refer to (Stelate et al., 2021).

The same imaging setup described above was used to obtain TIRFM images for the particle tracking and SRRF, but signals were recorded with Electron Multiplying Charge Coupled Device (EMCCD) camera (Andor iXon Ultra 897). For the excitation, 488 nm diode laser was used (300 mW; tuned to 0.5%) and 100 frames were acquired with 40 ms exposure time. NanoJ-SRRF plugin for ImageJ was used with default parameters (ring radius= 0.5, radial magnification=5, axes in ring=6).

### A-ESEM and TIRFM/A-ESEM CLEM

A-ESEM and CLEM were performed as described previously (Stelate et al., 2021). Briefly, samples were imaged using a patented (European patent number: 2195822) high-efficiency ionization secondary electron detector with an electrostatic separator (ISEDS), which allows low-dose and higher-resolution imaging (Neděla et al., 2018). For the A-ESEM, coverslips were placed on an aluminum stand and attached with double-sided tape for SEM. PM ghosts were imaged at a beam energy of 10 keV, a beam current of 30 pA, and a working distance of 8.5 mm. To prevent charging, the non-conductive samples were imaged at 200 Pa water vapor pressure in a vapourcore sample chamber. The software MAPS 2.5 (Thermo Fisher Scientific, Waltham, MA, USA) was used to correlate the A-ESEM and TIRF images. By maintaining the exact orientation of the coverslips in the sample chamber, we marked areas of the previously analyzed ghost images with the images from the CCD cameras in the A-ESEM. Then, the light microscope images previously acquired with a light microscope with 10× and 100× objectives were uploaded and used to assign the sample and locate the analyzed ghost images. Finally, the TIRFM images were uploaded and matched with the A-ESEM images. The exact superposition of all images was determined using the three-point alignment method based on rotation and resizing in two directions. The superimposed A-ESEM and TIRF images were used to correlate the position of gold nanoparticles whose positions and identities were detected using ISEDS.

### High-pressure SDS-FRL and TEM

2-day-old induced XVE-*Nt*PIN11T-GFP, XVE-*Nt*PIN2T-GFP and XVE-*Nt*PIN3bT-GFP cells were checked for their fluorescence and filtered through a 48 µm nylon mesh using a Nalgene filtration unit. Freeze fracturing and SDS-FRL protocol used here is based on the original protocol published for animal cells (Fujimoto, 1995). This protocol was modified for plant suspension-cultured cells as follows. Cells were transferred into gold-plated copper high-pressure freeze carriers with 100 µm cavity (Leica Microsystems GmbH, Wetzlar, Germany). The carrier assembly was inserted into the holder and immediately frozen in liquid nitrogen in the rapid freezing unit EM ICE (Leica, Germany). The frozen samples were transferred to liquid nitrogen and fractured in Freeze Fracture System Leica EM ACE900 at-140 °C, which split the PM into an E half (adjacent to the cell wall) and a P half (adjacent to the cytoplasm). Replicas of the P half were prepared by evaporating platinum (3 nm thick) from an electron beam gun at a 45° angle, followed by carbon coating (15 nm thick) at a 90° angle. Replicas were carefully immersed in PBS to separate them from the carrier and then transferred to 2.5% SDS (10mM Tris-HCl, pH 8.3) for 60 hours at room temperature with constant shaking on a rotary shaker at 3 rpm. After treatment with SDS, replicas were placed on 20 µl drops of blocking solution (3% BSA and 2% cold water fish skin gelatin in PBS) for 45 min at room temperature. After washing with 0.1% BSA in PBS for 10 min, replicas were placed on a 50 µl drop of X191 anti-GFP polyclonal rabbit primary antibody (A-11122, Thermofisher Scientific, Waltham, MA, USA) diluted 1:100 in 0.5% BSA in PBS. For positive control, replicas were stained with primary mouse monoclonal antibody against PM phosphoinositide PI(4,5)P2 (Z-A045, Echelon Biosciences Inc., Salt Lake City, UT, USA) diluted 1:400 in 0.5% BSA in PBS. Incubations were performed on parafilm in a humid chamber overnight at 4°C. After labeling, the replicas were washed three times with PBS and incubated for 2 hours at room temperature in 50 µl drops of goat anti-rabbit and goat anti-mouse secondary antibodies conjugated with colloidal 12 nm gold (111-205-144 and 115-205-075, JacksonImmuno Research Europe, Ltd, Ely, UK), both diluted 1:30 in PBTB 1% normal serum (Life Technologies Corp., Frederick, Maryland, USA) and 0.2% cold water fish skin gelatin (Merck KGaA, Darmstadt, Germany) in PBS. After immunogold labeling, the replicas were immediately rinsed three times in PBS and fixed with 0.5% glutaraldehyde (Thermo Fisher Scientific, MA, USA) in sodium boric acid buffer SB (Sörensen’s buffer, 0.1M Na/K phosphate buffer, pH 7.2-7.4) for 10 min at RT, washed five times with distilled water and imaged on formvar-coated grids. Electron microscopy was performed with a Jeol JEM 1400 flash operated at 80 Kv, equipped with Matataki Flash sCMOS camera (JEOL Ltd., Akishima, Tokyo, Japan).

### Protein extraction, isolation of PM fraction, co-IP and western blot

Induced 2-day-old BY-2 cells expressing *Nt*PIN11T-GFP, *Nt*PIN2T-GFP and *Nt*PIN3bT-GFP were checked microscopically for the presence of the fluorescence at the PM and harvested by filtration through 20 µm nylon mesh. 4 g of harvested cells were homogenised in liquid N_2_ using a pestle and mortar. The cell powder was resuspended on ice 1:1 (w:v) with lysis buffer (50mM PIPES, 2mM MgSO4, 2.5 mM EGTA, 300mM sucrose, 5mM DTT, pH 6.8), supplemented with a 1mM protease inhibitors cocktail (Sigma-Aldrich, P9599), 1mM PMSF and 5mM DTT and centrifuged at 4000 g for 10 min at 4°C. Pellet was discarded and the supernatant was centrifuged at 70000 g for 45 min at 4°C. To obtain a fraction with solubilized membrane proteins, the resulting pellet was resuspended in PM buffer (50 mM PIPES, 1 mM MgSO_4_, 1mM EGTA, pH 6.8) containing 1mM protease inhibitor cocktail and 40 mM CHAPS (Roche, 75621-03-3). The solution was shaken on ice for 50 min, centrifuged at 48 000 g for 60 min at 4°C and supernatant was collected, providing solubilized membrane fraction. The concentration of proteins in the solubilized membrane fraction was determined using the Bio-Rad Protein Assay (cat. no. 5000006). 1 ml of membrane protein fraction (1.5 mg proteins/ml) was mixed with 50 µl of super-paramagnetic beads (µMACS MicroBeads, GFP isolation kit, Miltenyi Biotec). The mixture was incubated on ice for 20 minutes and loaded into the column in a magnetic stand and GFP-tagged proteins bound to interacting proteins were isolated by washing with PM buffer without protease inhibitor cocktail, according to manufacturer protocol. Solubilized membrane fractions used as input to co-IP were separated by on 10% polyacrylamide SDS-PAGE gels and transferred to a nitrocellulose membrane by electroblotting (SemiDry, BioRad). Western blots were probed with rabbit anti-GFP (1:5000; cat. no. AS152987, Agrisera AB, Vännäs, Sweden). A chemiluminescence detection kit (Pierce™ ECL Western Blotting Substrate Thermo Scientific™ c.n 32109) with anti-rabbit secondary antibody coupled HRP (1:1000; Promega) was used to visualize immunolabeled proteins on the blot, and the chemiluminescence was recorded on an RTG film. The same procedure was used also for control cells expressing free GFP, which provided a set of candidates bound to GFP on membrane fractions. In addition, 35S::GFP cells were also used to prepare cytosolic fraction (isolated supernatant after the ultracentrifugation) to provide a set of candidates bound to free GFP in the cytosolic fraction.

### Preparation of co-IP protein fractions for MS analysis

Immunoprecipitaetd proteins were eluted by 100 µl of 100 mM Triethylammonium bicarbonate (TEAB) containing 2% sodium deoxycholate (SDC). Cysteines were subsequently reduced by boiling at 95°C for 10 min in 100mM TEAB containing 2% SDC, 40mM chloroacetamide, and 10mM Tris(2-carboxyethyl)phosphine (TCEP). Samples were further processed using single-pot solid-phase-enhanced sample preparation (SP3) method (Hughes et al., 2019). Briefly, 5 µl of SP3 beads was added to 30 µg of protein in lysis buffer and filled to 50 µl with 100mM TEAB. Protein binding was induced by the addition of ethanol to 60 % (v/v) final concentration. Samples were mixed and incubated for 5 min at RT. After binding, the tubes were placed into the magnetic rack and the unbound supernatant was discarded. Beads were subsequently washed two times with 180 µl of 80% ethanol. After washing, samples were digested with trypsin (1/30 trypsin/protein ratio) and reconstituted in 100mM TEAB at 37°C overnight. After digestion, samples were acidified with trifluoroacetic acid (TFA) and peptides were desalted using in-house made stage tips packed with C18 disks (Empore) according to (Rappsilber et al., 2007).

### nLC-MS^2^ analysis

Nano reversed-phase column (EASY-Spray column, 50 cm x 75 µm ID, PepMap C18, 2 µm particles, 100 Å pore size) was used for LC/MS analysis. Mobile phase buffer A was composed of water and 0.1% formic acid. Mobile phase B was composed of acetonitrile and 0.1% formic acid. Samples were loaded onto the trap column (Acclaim PepMap300, C18, 5 µm, 300 Å Wide Pore, 300 µm x 5 mm, 5 Cartridges) for 4 min at 15 μl/min. Loading buffer was composed of water, 2% acetonitrile, and 0.1% trifluoroacetic acid. Peptides were eluted with Mobile phase B gradient from 4% to 35% B in 60 min. Eluting peptide cations were converted to gas-phase ions by electrospray ionization and analyzed on a Thermo Orbitrap Fusion (Q-OT-qIT, Thermo Fisher). Survey scans of peptide precursors from 350 to 1400 m/z were performed at 120K resolution (at 200 m/z) with a 5 × 10^5^ ion count target. Tandem MS was performed by isolation at 1.5 Th with the quadrupole, HCD fragmentation with normalized collision energy of 30, and rapid scan MS analysis in the ion trap. The MS/MS ion count target was set to 10^4^ and the max injection time was 35 ms. Only those precursors with charge state 2-6 were sampled for MS/MS. The dynamic exclusion duration was set to 45 s with a 10 ppm tolerance around the selected precursor and its isotopes. Monoisotopic precursor selection was turned on. The instrument was run in top speed mode with 2 s cycles (Hebert et al., 2014).

All data were analyzed and quantified with the MaxQuant software 2.0.0.2 (Cox et al., 2014). The false discovery rate (FDR) was set to 1% for both proteins and peptides and we specified a minimum peptide length of seven amino acids. The Andromeda search engine was used for the MS/MS spectra search against the *Nicotiana tabacum* genome sequence database (Edwards et al., 2017). Enzyme specificity was set as C-terminal to Arg and Lys, also allowing cleavage at proline bonds and a maximum of two missed cleavages. Dithiomethylation of cysteine was selected as a fixed modification and N-terminal protein acetylation and methionine oxidation as variable modifications. The “match between runs” feature of MaxQuant was used to transfer identifications to other LC-MS/MS runs based on their masses and retention time (maximum deviation 0.7 min) and this was also used in quantification experiments. Quantifications were performed with the label-free algorithms described previously (Cox et al., 2014; O’Connell et al., 2018). Data analysis was performed using Perseus 1.6.15.0 software.

All proteins detected in membrane fractions that were also present in the cytosolic and membrane fractions from 35S::GFP cells were manually filtered out. The abundances of baits detected by peptides corresponding to GFP were increased 64 000-fold (35S::GFP), 1500-fold (Nitab4.5_0000812g0030.1; *Nt*PIN11T-GFP), 2500-fold (Nitab4.5_0000957g0120.1; *Nt*PIN2T-GFP) and 10 000-fold (Nitab4.5_0001832g0090.1; *Nt*PIN3bT-GFP) compared to the proteins with the lowest detected abundance. In all tested samples, peptides corresponding to GFP were identified using blast and GFP was confirmed to be present only in induced cells and in all control 35S::GFP cells. The stringent filtering against profiles free GFP expressing cells involved samples both from cytosolic and membrane fractions.

### Image analysis and statistics

TIRFM live cell images and PM ghost images of PMs were used to determine the coefficient of average intensity variation by generating a set of line profiles using Zen black (Carl Zeiss AG, Jena, Germany), which were analyzed in OriginPro by calculating the ratio of the standard deviation of fluorescence intensities along individual lines (STD) to the average intensities (AV) for each line profile, according to the following formula:

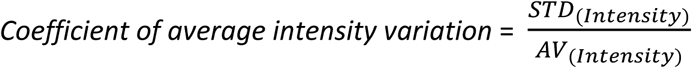

The size of the PM protein nanodomain in the TIRF and SRRF images was determined from the fluorescence intensity line generated across the nanodomains by Zen black software (Carl Zeiss AG, Jena, Germany). The FWHM was determined by manually selecting the peaks of the intensity profile using OriginPro 2017 (OriginLab Corporation, Northampton, MA, USA). A scatter box plot with error bars indicating the standard deviation was created using OriginPro. Single particle tracking and of mean square displacement estimation of nanodomains was performed on TIRFM images. The time series data were analyzed using SpatTrack as previously described (Lund & Wustner, 2013). The region of interest was cropped and subjected to particle tracking. Denoising was performed to improve particle detection (denoising = 0.5). The denoised image was generated by subtracting the boxcar-filtered image from the Gaussian blurred image (Crocker & Grier, 1996; Lund & Wustner, 2013). The threshold for particle detection was determined for each image using the “Find Threshold” function in SpatTrack. Particles were tracked for 50 frames (2 seconds) and trajectories were regenerated in SpatTrack. Only trajectories that lasted at least 400 ms (10 frames) were used for analysis, as this effectively reduced the amount of background noise features and false linkage and improved contrast resolution. The MSD was calculated based on the following equation:

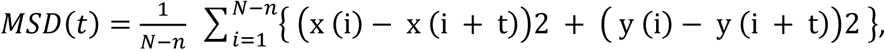

where N is the number of frames, x and y indicate the position of the particles, and n is the frame number corresponding to t (time). To determine the diffusion coefficient of the nanodomains, the MSD was fitted for each trajectory using the analytical Brownian diffusion model:

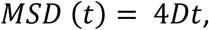

where D is the diffusion coefficient.

For spatial statistics of immunogold labeling patterns, we calculate the pair correlation function (PCF) using a software based on the previous report (Philimonenko et al., 2000), which could be freely downloaded from <u>nucleus.img.cas.cz/gold/program/IE/5a.htm</u>. This software quantifies the number of gold particles at a given distance (d) between r’ and r” (ring shape), normalized by the area of each ring and the average particle density, and averages over all particles. Therefore, if a protein population is randomly distributed, the pair correlation function describing its organization approaches 1. If the proteins are not randomly distributed and exhibit clustering or aggregation, PCF is greater than 1, and the peak of the shortest histogram and greater than 1 correspond to the size of the cluster. The correlation function of the pairs under these conditions is given by the following equation:

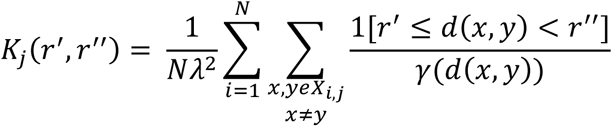

For the majority of plots, the R-based online tool PlotsOfData was used to depict actual data together with their summaries (Postma & Goedhart, 2019).

### Molecular dynamics simulation

Amino acid sequences of GFP-tagged versions of *Nt*PIN11T, *Nt*PIN2T and *Nt*PIN3bT (Müller et al., 2019) were used for AlphaFold *in silico* protein folding using Google Colab platform (Jumper et al., 2021; Mirdita et al., 2022). PIN2T, PIN3bT and PIN11T homodimers were predicted using the Colabfold (version 1.5.2) implementation of AlphaFold2-Multimer v3 (Mirdita et al., 2022). The simulations were performed using the GROMACS software (Abraham et al., 2015) and MARTINI 2.2 coarse-grained (CG) force-field (De Jong et al., 2013). Predicted PIN2T, PIN3bT and PIN2T protein structures were converted to CG representation using CHARMM-GUI web server (Jo et al., 2008; Wu et al., 2014). Loop regions (PIN2T210- 453, PIN3bT210-455 and PIN11T210-427) were excluded from the simulations. Elastic Network in Dynamics (ElNeDyn) was applied to retain the secondary and tertiary structure (Periole et al., 2009). Protein in CG representation was inserted in 15×15 nm membrane composed of 50 % PC, 50 % PE in the upper leaflet and 36 % PC 36 % PE 10 % PS 10 % PA 5 % PI4P and 3 % PI(4,5)P2 in the lower leaflet using insane.py script (Wassenaar et al., 2015). The charge of PA was modified to-2. Protein membrane system was inserted in 15×15×10 nm simulation box, solvated with 0.15 M NaCl in water and neutralized. The system was energy minimized using the steepest descent algorithm up to the maximum of 5000 steps and equilibrated for 4750 ps. Position restraints on protein backbone and side chains were progressively released during equilibration. Production run was performed in the NPT ensemble. Pressure was set to 1 bar and maintained with the Parrinello-Rahman barostat, with a coupling constant of 12.0 ps and compressibility 3.4e-4. Temperature was set to 323 K and maintained with the v-rescale thermostat with a coupling constant of 1.0 ps. The bond lengths were constrained using the LINCS algorithm. Van der Waals and Coulomb cut-offs were set to 1.1 nm. Timestep was set to 20 fs. In total 5 production runs, 5 μs each starting from different initial velocities were performed. PI(4,5)P2 clustering was analysed using GROMACS in-build tool gmx densmap over the last 3 μs. To track protein diffusion in the membrane, X and Y positions of centre of mass of the protein were tracked using VMD software (Humphrey et al., 1996) over the whole trajectory.

## Acknowledgments

This work was supported by the Czech Science Foundation, projects n. 22-25799S (V.N., J.P.) and 22-35680M (R.P.) and Grant Agency of Charles University, project n. 1374120 (A.S.). We acknowledge a helpful discussions on the image analysis of Dominik Pinkas. Microscopy was performed in Core facility Microscopy Centre, Electron Microscopy Facility, Institute of Molecular Genetics and Imaging Facility of the Institute of Experimental Botany AS CR, both supported by the MEYS CR (LM2023050 Czech-BioImaging). Computational resources used for structural modeling were provided by the e-INFRA CZ project (ID:90254), supported by the Ministry of Education, Youth and Sports of the Czech Republic.

## References

Abraham, M.J., Murtola, T., Schulz, R., Páll, S., Smith, J.C., Hess, B., & Lindah, E. (2015). Gromacs: High performance molecular simulations through multi-level parallelism from laptops to supercomputers. SoftwareX, 1-2, 19/25.

Adamowski, M., & Friml, J. (2015). PIN-dependent auxin transport: Action, regulation, and evolution. Plant Cell, 27(1), 20-32.

Band, L. R., Wells, D. M., Fozard, J. A., Ghetiu, T., French, A. P., Pound, M. P., Wilson, M. H., Yu, L., Li, W., Hijazi, H.I., Oh, J., Pearce, S. P., Perez-Amador, M.A., Yun, J., Kramer, E., Alonso, J.M., Godin, C., Vernoux, T., Hodgman, T.C., Pridmore, T.P., Swarup, R., King, J.R., & Bennett, M.J. (2014). Systems Analysis of Auxin Transport in the *Arabidopsis* Root Apex. The Plant Cell, 26(3), 862-875.

Bassukas, A. E. L., Xiao, Y., & Schwechheimer, C. (2022). Phosphorylation control of PIN auxin transporters. Current Opinion in Plant Biology, 65, 102146.

Bücherl, C. A., Jarsch, I.K., Schudoma, C., Segonzac, C., Mbengue, M., Robatzek, S., MacLean, D., Ott, T., & Zipfel, C. (2017). Plant immune and growth receptors share common signalling components but localise to distinct plasma membrane nanodomains. ELife, 6.

Busto, J. V, Elting, A., Haase, D., Spira, F., Kuhlman, J., Schäfer-Herte, M., & Wedlich-Söldner, R. (2018). Lateral plasma membrane compartmentalization links protein function and turnover. The EMBO Journal, 37(16).

Corradi, V., Mendez-Villuendas, E., Ingólfsson, H., Gu, Ruo, X., Siuda, I., Melo, M.N., Moussatova, A., Degagné, L.J., Sejdiu, B.I., Singh, G., Wassenaar, T.A., Magnero, K.D., Marrink, S.J., & Tieleman, D.P. (2018). Lipid-Protein Interactions Are Unique Fingerprints for Membrane Proteins. ACS Central Science, 4, 709-717.

Cox, J., Hein, M. Y., Luber, C.A., Paron, I., Nagaraj, N., & Mann, M. (2014). Accurate Proteome-wide Label-free Quantification by Delayed Normalization and Maximal Peptide Ratio Extraction, Termed MaxLFQ. Molecular & Cellular Proteomics, 13(9), 2513-2526.

Crocker, J.C., & Grier, D.G. (1996). Methods of Digital Video Microscopy for Colloidal Studies. Journal of Colloid and Interface Science, 179(1), 298-310.

De Jong D.H., Singh, G., Bennett, W.F.D., Arnarez, C., Wassenaar, T.A., Schäfer, L.V., Periole, X., Tieleman, D.P., & Marrink, S.J. (2013). Improved parameters for the martini coarse-grained protein force field. Journal of Chemical Theory and Computation 9, 687-697.

Delbarre, A., Muller, P., Imhoff, V., & Guern, J. (1996). Comparison of mechanisms controlling uptake and accumulation of 2,4-dichlorophenoxy acetic acid, naphthalene-1-acetic acid, and indole-3- acetic acid in suspension-cultured tobacco cells. Planta, 198(4), 532-541.

Dubey, S. M., Serre, N. B. C., Oulehlová, D., Vittal, P., & Fendrych, M. (2021). No Time for Transcription-Rapid Auxin Responses in Plants. Cold Spring Harbor Perspectives in Biology, 13(8), a039891.

Edwards, K. D., Fernandez-Pozo, N., Drake-Stowe, K., Humphry, M., Evans, A.D., Bombarely, A., Allen, F., Hurst, R., White, B., Kernodle, S. P., Bromley, J. R., Sanchez-Tamburrino, J. P., Lewis, R. S., & Mueller, L.A. (2017). A reference genome for Nicotiana tabacum enables map-based cloning of homeologous loci implicated in nitrogen utilization efficiency. BMC Genomics, 18(1), 448.

Feraru, E., Feraru, M. I., Kleine-Vehn, J., Martinière, A., Mouille, G., Vanneste, S., Vernhettes, S., Runions, J., & Friml, J. (2011). PIN polarity maintenance by the cell wall in Arabidopsis. Current Biology, 21(4), 338-343.

Friml, J. (2022). Fourteen Stations of Auxin. Cold Spring Harbor Perspectives in Biology, 14(5), a039859.

Fujimoto, K. (1995). Freeze-fracture replica electron microscopy combined with sds digestion for cytochemical labeling of integral membrane proteins: Application to the immunogold labeling of intercellular junctional complexes. Journal of Cell Science, 108(11), 3443–3449.

Gadeyne, A., Sánchez-Rodríguez, C., Vanneste, S., Di Rubbo, S., Zauber, H., Vanneste, K., Van Leene, J., De Winne, N., Eeckhout, D., Persiau, G., Van De Slijke, E., Cannoot, B., Vercruysse, L., Mayers, J.R., Adamowski, M., Kania, U., Ehrlich, M., Schweighofer, A., Ketelaar, T., Maere, S., Bednarek, S.Y., Friml. J., Gevaert, K, Witters, E., Russinova, E., Persson, S., De Jaeger, G., & Van Damme, D. (2014). The TPLATE Adaptor Complex Drives Clathrin-Mediated Endocytosis in Plants. Cell, 156(4), 691-704.

Glanc, M., Fendrych, M., & Friml, J. (2019). PIN2 Polarity Establishment in Arabidopsis in the Absence of an Intact Cytoskeleton. Biomolecules, 9(6), 222.

Gustafsson, N., Culley, S., Ashdown, G., Owen, D. M., Pereira, P. M., & Henriques, R. (2016). Fast live-cell conventional fluorophore nanoscopy with ImageJ through super-resolution radial fluctuations. Nature Communications, 7(1), 12471.

Hebert, A. S., Richards, A. L., Bailey, D. J., Ulbrich, A., Coughlin, E. E., Westphall, M. S., & Coon, J. J. (2014). The One Hour Yeast Proteome. Molecular & Cellular Proteomics, 13(1), 339-347.

Humphrey, W., Dalke, A., Schulten, K. (1996). VMD: Visual molecular dynamics. Journal of Molecular Graphics 14, 33-38.

Ivanchenko, M. G., Zhu, J., Wang, B., Medvecká, E., Du, Y., Azzarello, E., Mancuso, S., Megraw, M., Filichkin, S., Dubrovsky, J. G., Friml, J., & Geisler, M. (2015). The cyclophilin A DIAGEOTROPICA gene affects auxin transport in both root and shoot to control lateral root formation. Development.

Jacobson, K., Liu, P., & Lagerholm, B. C. (2019). The Lateral Organization and Mobility of Plasma Membrane Components. Cell, 177(4), 806-819.

Jo, S., Kim, T., Iyer, V.G., Im, W. (2008). CHARMM-GUI: A web-based graphical user interface for CHARMM. Journal of Computational Chemistry 29, 1859-1865.

Jones, A. M., Xuan, Y., Xu, M., Wang, R. S., Ho, C. H., Lalonde, S., You, C. H., Sardi, M. I., Parsa, S. A., Smith-Valle, E., Su, T., Frazer, K. A., Pilot, G., Pratelli, R., Grossmann, G., Acharya, B. R., Hu, H. C., Engineer, C., Villiers, F., Ju, C., Takeda, K., Su, Z., Dong, Q., Assmann, S.M., Chen, J., Kwak, J.M., Schroeder, J.I., Albert, R., Rhee, S.Y., & Frommer, W. B. (2014). Border control - A membrane-linked interactome of Arabidopsis. Science, 344(6185), 711-716.

Jumper, J., Evans, R., Pritzel, A., Green, T., Figurnov, M., Ronneberger, O., Tunyasuvunakool, K., Bates, R., Žídek, A., Potapenko, A., Bridgland, A., Meyer, C., Kohl, S. A. A., Ballard, A. J., Cowie, A., Romera-Paredes, B., Nikolov, S., Jain, R., Adler, J., Back, T., Petersen, S., Reiman, D., Clancy, E., Zielinski, M., Steinegger, M., Pacholska, M., Berghammer, T., Bodenstein, S., Silver, D., Vinyals, O., Senior, A.W., Kavukcuoglu, K., Kohli, P., & Hassabis, D. (2021). Highly accurate protein structure prediction with AlphaFold. Nature, 596(7873), 583-589.

Kashkan, I., Hrtyan, M., Retzer, K., Humpolíčková, J., Jayasree, A., Filepová, R., Vondráková, Z., Simon, S., Rombaut, D., Jacobs, T. B., Frilander, M. J., Hejátko, J., Friml, J., Petrášek, J., & Růžička, K. (2022). Mutually opposing activity of PIN7 splicing isoforms is required for auxin-mediated tropic responses in Arabidopsis thaliana. New Phytologist, 233(1), 329-343.

Ke, M., Ma, Z., Wang, D., Sun, Y., Wen, C., Huang, D., Chen, Z., Yang, L., Tan, S., Li, R., Friml, J., Miao, Y., & Chen, X. (2021). Salicylic acid regulates *PIN2* auxin transporter hyperclustering and root gravitropic growth via *Remorin*-dependent lipid nanodomain organisation in *Arabidopsis thaliana*. New Phytologist, 229(2), 963-978.

Kimura, S., Laosinchai, W., Itoh, T., Cui, X., Linder, C.R., & Malcolm Brown, R. (1999). Immunogold labeling of rosette terminal cellulose-synthesizing complexes in the vascular plant Vigna angularis. Plant Cell, 11(11), 2075-2085.

Kitakura, S., Vanneste, S., Robert, S., Löfke, C., Teichmann, T., Tanaka, H., & Friml, J. (2011). Clathrin mediates endocytosis and polar distribution of PIN auxin transporters in Arabidopsis. Plant Cell, 23(5), 1920-1931.

Kleine-Vehn, J., Łangowski, Ł., Wiśniewska, J., Dhonukshe, P., Brewer, P. B., & Friml, J. (2008). Cellular and Molecular Requirements for Polar PIN Targeting and Transcytosis in Plants. Molecular Plant, 1(6), 1056-1066.

Kleine-Vehn, J., Wabnik, K., Martinière, A., Łangowski, Ł., Willig, K., Naramoto, S., Leitner, J., Tanaka, H., Jakobs, S., Robert, S., Luschnig, C., Govaerts, W., W Hell, S., Runions, J., & Friml, J. (2011). Recycling, clustering, and endocytosis jointly maintain PIN auxin carrier polarity at the plasma membrane. Molecular Systems Biology, 7(540), 1-13.

Korbei, B., Moulinier-Anzola, J., De-Araujo, L., Lucyshyn, D., Retzer, K., Khan, M. A., & Luschnig, C. (2013). Arabidopsis TOL proteins act as gatekeepers for vacuolar sorting of PIN2 plasma membrane protein. Current Biology, 23(24), 2500-2505.

Kramer, E. M., Rutschow, H. L., & Mabie, S. S. (2011). AuxV: a database of auxin transport velocities. Trends in Plant Science, 16(9), 461-463.

Krtková, J., Zimmermann, A., Schwarzerová, K., & Nick, P. (2012). Hsp90 binds microtubules and is involved in the reorganization of the microtubular network in angiosperms. Journal of Plant Physiology, 169(14), 1329-1339.

Kuthanová, A., Fischer, L., Nick, P., & Opatrný, Z. (2008). Cell cycle phase-specific death response of tobacco BY-2 cell line to cadmium treatment. *Plant*, Cell & Environment, 31(11), 1634-1643.

Laňková, M., Humpolíčková, J., Vosolsobě, S., Cit, Z., Lacek, J., Čovan, M., Čovanová, M., Hof, M., & Petrášek, J. (2016). Determination of Dynamics of Plant Plasma Membrane Proteins with Fluorescence Recovery and Raster Image Correlation Spectroscopy. Microscopy and Microanalysis, 22(02), 290-299.

Leitner J., Retzer, K., Malenica, N., Bartkeviciute, R., Lucyshyn, D., Jäger, G., Korbei, B., Byström, A., & Luschnig, C. (2015). Meta-regulation of Arabidopsis Auxin Responses Depends on tRNA Maturation. Cell Reports 11, 516-526.

Li, H., von Wangenheim, D., Zhang, X., Tan, S., Darwish-Miranda, N., Naramoto, S., Wabnik, K., De Rycke, R., Kaufmann, W. A., Gütl, D., Tejos, R., Grones, P., Ke, M., Chen, X., Dettmer, J., & Friml, J. (2021). Cellular requirements for PIN polar cargo clustering in *Arabidopsis thaliana*. New Phytologist, 229(1), 351-369.

Liu, E., MacMillan, C. P., Shafee, T., Ma, Y., Ratcliffe, J., van de Meene, A., Bacic, A., Humphries, J., & Johnson, K. L. (2020). Fasciclin-Like Arabinogalactan-Protein 16 (FLA16) Is Required for Stem Development in Arabidopsis. Frontiers in Plant Science, 11.

Lund, F.W., & Wustner, D. (2013). A comparison of single particle tracking and temporal image correlation spectroscopy for quantitative analysis of endosome motility. Journal of Microscopy, 252(2), 169-188.

Ma, Y., MacMillan, C. P., de Vries, L., Mansfield, S. D., Hao, P., Ratcliffe, J., Bacic, A., & Johnson, K. L. (2022). FLA11 and FLA12 glycoproteins fine-tune stem secondary wall properties in response to mechanical stresses. The New Phytologist, 233(4), 1750-1767.

Marhava, P. (2022). Recent developments in the understanding of PIN polarity. New Phytologist, 233(2), 624-630.

Martinez, C.C., Koenig, D., Chitwood, D.H., & Sinha, N.R. (2016). A sister of PIN1 gene in tomato (Solanum lycopersicum) defines leaf and flower organ initiation patterns by maintaining epidermal auxin flux. Developmental Biology, 419(1), 85-98.

Martinière, A., Lavagi, I., Nageswaran, G., Rolfe, D. J., Maneta-Peyret, L., Luu, D.-T., Botchway, S. W., Webb, S. E. D., Mongrand, S., Maurel, C., Martin-Fernandez, M. L., Kleine-Vehn, J., Friml, J., Moreau, P., & Runions, J. (2012). Cell wall constrains lateral diffusion of plant plasma-membrane proteins. Proceedings of the National Academy of Sciences of the United States of America, 109(31), 12805-12810.

Martiniere, A., Lavagi, I., Nageswaran, G., Rolfe, D. J., Maneta-Peyret, L., Luu, D.-T., Botchway, S. W., Webb, S. E. D., Mongrand, S., Maurel, C., Martin-Fernandez, M. L., Kleine-Vehn, J., Friml, J., Moreau, P., & Runions, J. (2012). Cell wall constrains lateral diffusion of plant plasma-membrane proteins. Proceedings of the National Academy of Sciences, 109(31), 12805-12810.

Martinière, A., & Runions, J. (2013). Protein diffusion in plant cell plasma membranes: the cell-wall corral. Frontiers in Plant Science, 4, 515.

Martinière, A., & Zelazny, E. (2021). Membrane nanodomains and transport functions in plant. Plant Physiology, 1-17.

McKenna, J. F., Rolfe, D. J., Webb, S. E. D., Tolmie, A. F., Botchway, S. W., Martin-Fernandez, M. L., Hawes, C., & Runions, J. (2019). The cell wall regulates dynamics and size of plasma-membrane nanodomains in Arabidopsis. Proceedings of the National Academy of Sciences of the United States of America, 116(26), 12857-12862.

McWhite, C.D., Papoulas, O., Drew, K., Cox, R.M., June, V., Dong, O. X., Kwon, T., Wan, C., Salmi, M.L., Roux, S.J., Browning, K.S., Chen, Z.J., Ronald, P.C., & Marcotte, E.M. (2020). A Pan-plant Protein Complex Map Reveals Deep Conservation and Novel Assemblies. Cell, 181(2), 460-474.e14.

Mellor, N. L., Voß, U., Ware, A., Janes, G., Barrack, D., Bishopp, A., Bennett, M.J., Geisler, M., Wells, D. M., & Band, L. R. (2022). Systems approaches reveal that ABCB and PIN proteins mediate co-dependent auxin efflux. The Plant Cell, 34(6), 2309-2327.

Mirdita, M., Schütze, K., Moriwaki, Y., Heo, L., Ovchinnikov, S., & Steinegger, M. (2022). ColabFold: making protein folding accessible to all. Nature Methods, 19(6), 679-682.

Mravec, J., Petrášek, J., Li, N., Boeren, S., Karlova, R., Kitakura, S., Pařezová, M., Naramoto, S., Nodzyński, T., Dhonukshe, P., Bednarek, S. Y., Zažímalová, E., De Vries, S., & Friml, J. (2011). Cell plate restricted association of DRP1A and PIN proteins is required for cell polarity establishment in arabidopsis. Current Biology, 21(12), 1055–1060. 10.1016/j.cub.2011.05.018

Müller, K., Dobrev, P. I., Pěnčík, A., Hošek, P., Vondráková, Z., Filepová, R., Malínská, K., Brunoni, F., Helusová, L., Moravec, T., Retzer, K., Harant, K., Novák, O., Hoyerová, K., & Petrášek, J. (2021). DIOXYGENASE FOR AUXIN OXIDATION 1 catalyzes the oxidation of IAA amino acid conjugates. Plant Physiology, 187(1), 103-115.

Müller, K., Hošek, P., Laňková, M., Vosolsobě, S., Malínská, K., Čarná, M., Fílová, M., Dobrev, P. I., Helusová, M., Hoyerová, K., & Petrášek, J. (2019). Transcription of specific auxin efflux and influx carriers drives auxin homeostasis in tobacco cells. Plant Journal, 100(3), 627-640.

Müller, K., Dobrev, P.I., Pěnčík, A., Hošek, P., Vondráková, Z., Filepová, R., Malínská, K., Brunoni, F., Helusová, L., Moravec, T., Retzer, K., Harant, K., Novák, O., Hoyerová, K., & Petrášek, J. (2020). DIOXYGENASE FOR AUXIN OXIDATION 1 catalyzes the oxidation of IAA amino acid conjugates. Plant Physiology 187, 103-115.

Nagata, T., Nemoto, Y., & Hasezawa, S. (1992). Tobacco BY-2 Cell Line as the “HeLa” Cell in the Cell Biology of Higher Plants. International Review of Cytology, 132(C), 1-30.

Narasimhan, M., Gallei, M., Tan, S., Johnson, A., Verstraeten, I., Li, L., Rodriguez, L., Han, H., Himschoot, E., Wang, R., Vanneste, S., Sánchez-Simarro, J., Aniento, F., Adamowski, M., & Friml, J. (2021). Systematic analysis of specific and nonspecific auxin effects on endocytosis and trafficking. Plant Physiology, 186(2), 1122-1142.

Narasimhan, M., Johnson, A., Prizak, R., Kaufmann, W. A., Tan, S., Casillas-Pérez, B., & Friml, J. (2020). Evolutionarily unique mechanistic framework of clathrin-mediated endocytosis in plants. ELife, 9, 1-30.

Neubergerová, M., Pleskot, R. (2024). Plant protein-lipid interfaces studied by molecular dynamics simulations. Journal of Experimental Botany, 75, 5237-5250.

Neděla, V., Tihlaříková, E., Runštuk, J., & Hudec, J. (2018). High-efficiency detector of secondary and backscattered electrons for low-dose imaging in the ESEM. Ultramicroscopy, 184, 1-11.

O’Connell, J. D., Paulo, J. A., O’Brien, J.J., & Gygi, S.P. (2018). Proteome-Wide Evaluation of Two Common Protein Quantification Methods. Journal of Proteome Research, 17(5), 1934-1942.

O’Connor, D. L., Elton, S., Ticchiarelli, F., Hsia, M. M., Vogel, J.P., & Leyser, O. (2017). Cross-species functional diversity within the PIN auxin efflux protein family. ELife, 6.

Petrášek, J., Černá, A., Schwarzerová, K., Elčkner, M., Morris, D. A., & Zažímalová, E. (2003). Do Phytotropins Inhibit Auxin Efflux by Impairing Vesicle Traffic? Plant Physiology, 131(1), 254–263.

Petrášek, J., & Zažímalová, E. (2006). The BY-2 Cell Line as a Tool to Study Auxin Transport. Biotechnology in Agriculture and Forestry, 58, 107–115. 10.1007/3-540-32674-X_8

Philimonenko, A. A., Janáček, J., & Hozák, P. (2000). Statistical Evaluation of Colocalization Patterns in Immunogold Labeling Experiments. Journal of Structural Biology, 132(3), 201-210.

Periole, X., Cavalli, M., Marrink, S.J., & Ceruso, M.A. (2009). Combining an elastic network with a coarse-grained molecular force field: Structure, dynamics, and intermolecular recognition. Journal of Chemical Theory and Computation, 5, 2531-2543.

Platre, M. P., Bayle, V., Armengot, L., Bareille, J., del Mar Marquès-Bueno, M., Creff, A., Maneta-Peyret, L., Fiche, J. B., Nollmann, M., Miège, C., Moreau, P., Martinière, A., & Jaillais, Y. (2019). Developmental control of plant Rho GTPase nano-organization by the lipid phosphatidylserine. Science, 364(6435), 57–62.

Postma, M., & Goedhart, J. (2019). Plots Of Data-A web app for visualizing data together with their summaries. PLOS Biology, 17(3), e3000202.

Rappsilber, J., Mann, M., & Ishihama, Y. (2007). Protocol for micro-purification, enrichment, pre-fractionation and storage of peptides for proteomics using Stage Tips. Nature Protocols, 2(8), 1896-1906.

Sarrion-Perdigones, A., Vazquez-Vilar, M., Palaci, J., Castelijns, B., Forment, J., Ziarsolo, P., Blanca, J., Granell, A., & Orzaez, D. (2013). GoldenBraid 2.0: A Comprehensive DNA Assembly Framework for Plant Synthetic Biology. Plant Physiology, 162(3), 1618–1631.

Sonobe, S., & Takahashi, S. (1994). Association of Microtubules with the Plasma Membrane of Tobacco BY-2 Cells in Vitro. Plant and Cell Physiology, 35(3), 451-460.

Stelate, A., Tihlaříková, E., Schwarzerová, K., Neděla, V., & Petrášek, J. (2021). Correlative Light-Environmental Scanning Electron Microscopy of Plasma Membrane Efflux Carriers of Plant Hormone Auxin. Biomolecules, 11(10), 1407.

Su, N., Zhu, A., Tao, X., Ding, Z. J., Chang, S., Ye, F., Zhang, Y., Zhao, C., Chen, Q., Wang, J., Zhou, C. Y., Guo, Y., Jiao, S., Zhang, S., Wen, H., Ma, L., Ye, S., Zheng, S. J., Yang, F., Wu, S., & Guo, J. (2022). Structures and mechanisms of the Arabidopsis auxin transporter PIN3. Nature, 1-2.

Tanaka, K., Hayashi, K., Natsume, M., Kamiya, Y., Sakakibara, H., Kawaide, H., & Kasahara, H. (2014). UGT74D1 Catalyzes the Glucosylation of 2-Oxindole-3-Acetic Acid in the Auxin Metabolic Pathway in Arabidopsis. Plant and Cell Physiology, 55(1), 218-228.

Tateno, M., Brabham, C., & DeBolt, S. (2016). Cellulose biosynthesis inhibitors - a multifunctional toolbox. Journal of Experimental Botany, 67(2), 533-542.

Titapiwatanakun, B., Blakeslee, J. J., Bandyopadhyay, A., Yang, H., Mravec, J., Sauer, M., Cheng, Y., Adamec, J., Nagashima, A., Geisler, M., Sakai, T., Friml, J., Peer, W. A., & Murphy, A. S. (2009). ABCB19/PGP19 stabilises PIN1 in membrane microdomains in Arabidopsis. Plant Journal, 57(1), 27-44.

Tolmie, F., Poulet, A., McKenna, J., Sassmann, S., Graumann, K., Deeks, M., & Runions, J. (2017). The cell wall of Arabidopsis thaliana influences actin network dynamics. Journal of Experimental Botany, 68(16), 4517-4527.

Ung, K. L., Winkler, M., Schulz, L., Kolb, M., Janacek, D. P., Dedic, E., Stokes, D. L., Hammes, U. Z., & Pedersen, B. P. (2022). Structures and mechanism of the plant PIN-FORMED auxin transporter. Nature, 609(7927), 605-610.

Vanneste, S., & Friml, J. (2009). Auxin: A Trigger for Change in Plant Development. Cell, 136(6), 1005-1016.

Vieten, A., Vanneste, S., Wiśniewska, J., Benková, E., Benjamins, R., Beeckman, T., Luschnig, C., & Friml, J. (2005). Functional redundancy of PIN proteins is accompanied by auxin-dependent cross-regulation of PIN expression. Development, 132(20), 4521-4531.

Wassenaar, T.A., Ingólfsson, H.I., Böckmann, R.A., Tieleman, D.P., Marrink, S.J. (2015). Computational lipidomics with insane: A versatile tool for generating custom membranes for molecular simulations. Journal of Chemical Theory and Computation, 11, 2144-2155.

Wiśniewska, J., Xu, J., Seifertová, D., Brewer, P. B., Růžička, K., Blilou, I., Rouquié, D., Benková, E., Scheres, B., & Friml, J. (2006). Polar PIN Localization Directs Auxin Flow in Plants. Science, 312(5775), 883-883.

Wu, E.L., Cheng, X., Jo, S., Rui, H., Song, K.C., Dávila-Contreras, E.M., Qi, Y., Lee, J., Monje-Galvan, V., Venable, R.M., Klauda, J.B., & Wonpil I. (2014). CHARMM-GUI Membrane Builder toward realistic biological membrane simulations. Journal of Computational Chemistry 35, 1997-2004.

Xi, W., Gong, X., Yang, Q., Yu, H., & Liou, Y.C. (2016). Pin1At regulates PIN1 polar localization and root gravitropism. Nature Communications, 7(1), 10430.

Yang, Z., Xia, J., Hong, J., Zhang, C., Wei, H., Ying, W., Sun, C., Sun, L., Mao, Y., Gao, Y., Tan, S., Friml, J., Li, D., Liu, X., & Sun, L. (2022). Structural insights into auxin recognition and efflux by Arabidopsis PIN1. Nature 609, 611-615.

Zuo, J., Niu, Q.-W., & Chua, N.H. (2000). An estrogen receptor-based transactivator XVE mediates highly inducible gene expression in transgenic plants. Plant Journal, 24(2), 265-273.

