## Supplemental files for "Nanodomain distribution and function of PIN-FORMED auxin efflux carriers in the plasma membrane of tobacco cells are defined by their interactions with the cell wall"

#### Supplementary Information

### Nano-organization and function of tobacco PINFORMED auxin efflux carriers are defined by their specific interactions with the cell wall

Ayoub Stelate, Kateřina Malínská, Eva Tihlaříková, Karel Müller, Roberta Vaculíková, Martina Knirsch, Zuzana Vondráková, Erik Vlčák, Katarzyna Retzer, Michaela Neubergerová, Roman Pleskot, Karel Harant, Vlada Filimonenko, Kateřina Schwarzerová, Vilém Neděla, and Jan Petrášek*

*Correspondence: Jan Petrášek


**This PDF file includes:**

SI Figures S1-S5
SI Tables S1-S4
SI References

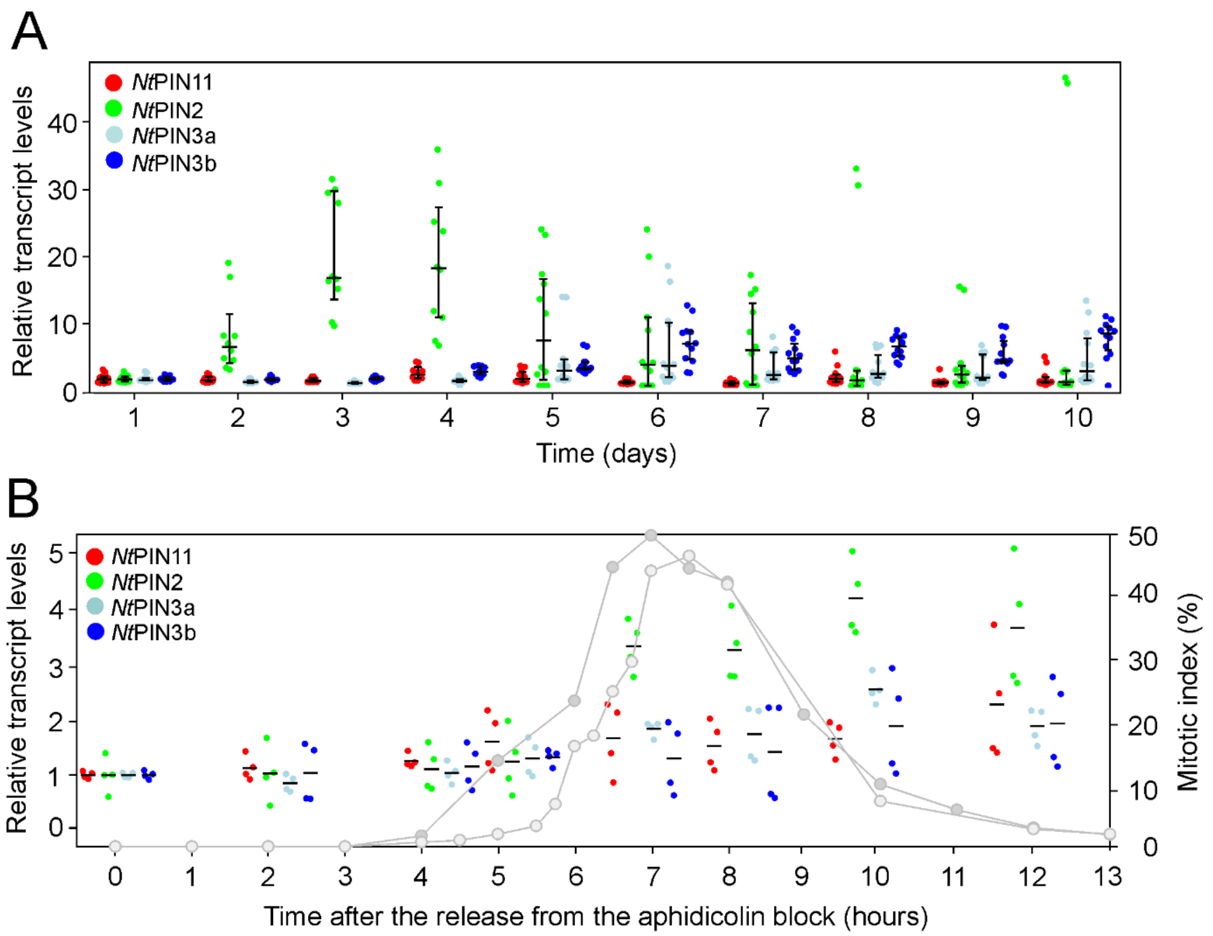


**SI Fig. S1: RT-qPCR transcriptional profiling of tobacco *Nt*PINs. (A)** Relative transcript levels during the life cycle of *NtPINs* in wild-type tobacco BY-2 cells. Values for days 2-10 represent fold changes in relation to day 1 (24 hours after the inoculation). The number of sample points from at least three biological repetitions and three technical repetitions in each of them for days 1 and 5-10 is 12, for days 2-4 is 10. Horizontal lines indicate the median. For each median, the vertical bar indicates the 95% confidence interval determined by bootstrapping. **(B)** Relative transcript levels during the cell cycle of *NtPINs* in wild-type tobacco BY-2 cells. Values for 0-13 hours after the release from the aphidicolin block represent fold changes in relation to time 0 (time of the release from the aphidicolin block). The number of sample points is four from two independent synchronizations and two technical repetitions each. Horizontal lines indicate the mean. Grey lines show the mitotic index for two biological repetitions. The color coding is indicated in the legend above the plot.

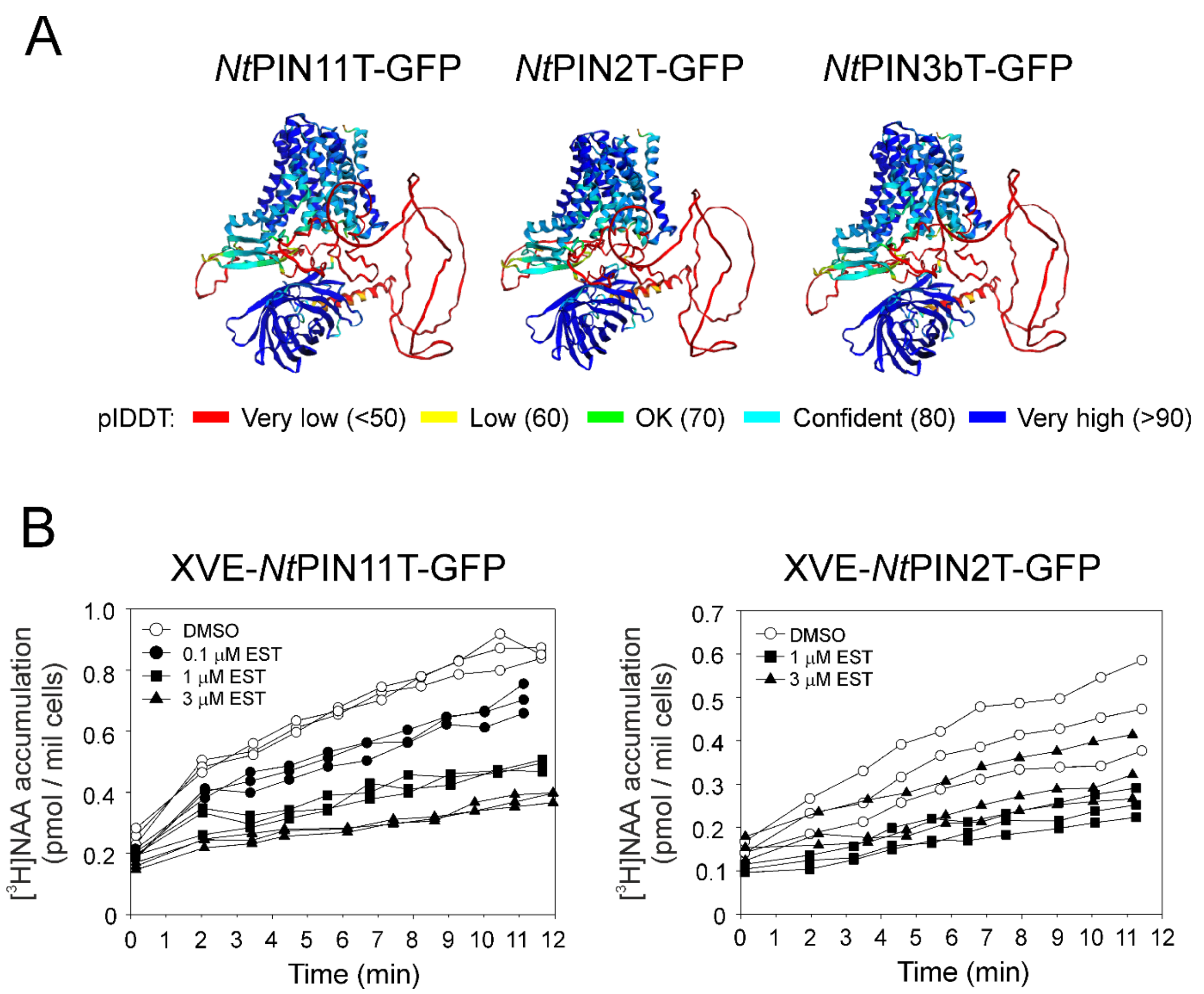


**SI Fig. S2: Dose-dependent auxin efflux activity and structure of GFP-tagged *Nt*PINs. (A)** AlphaFold2 predictions of the protein structures of *Nt*PIN11T-GFP, *Nt*PIN2T-GFP and *Nt*PIN3bT-GFP. GFP is inserted into the cytosolic loop. Per-residue estimate of the model's predicted score is shown below the image. Note that both the GFP and integral plasma membrane (PM) domains of tobacco PINs are predicted with a very high degree of prediction accuracy. **(B)** β-estradiol dose-dependent [$^3$H]NAA accumulation kinetics in XVE-*Nt*PIN11T-GFP and XVE-*Nt*PIN2T-GFP cells. XVE-*Nt*PIN11T-GFP cells induced for 48 h with 0, 0.1, 1 and 3 μM β-estradiol showing a clear dose-dependent decrease in the [$^3$H]NAA accumulations, reflecting the activity of the auxin efflux carrier (left plot). XVE-*Nt*PIN2T-GFP cells induced for 48 h with 0, 1 and 3 μM β-estradiol showing a dose-dependent decrease in the [$^3$H]NAA accumulation, reflecting the activity of the auxin efflux carrier (right plot).

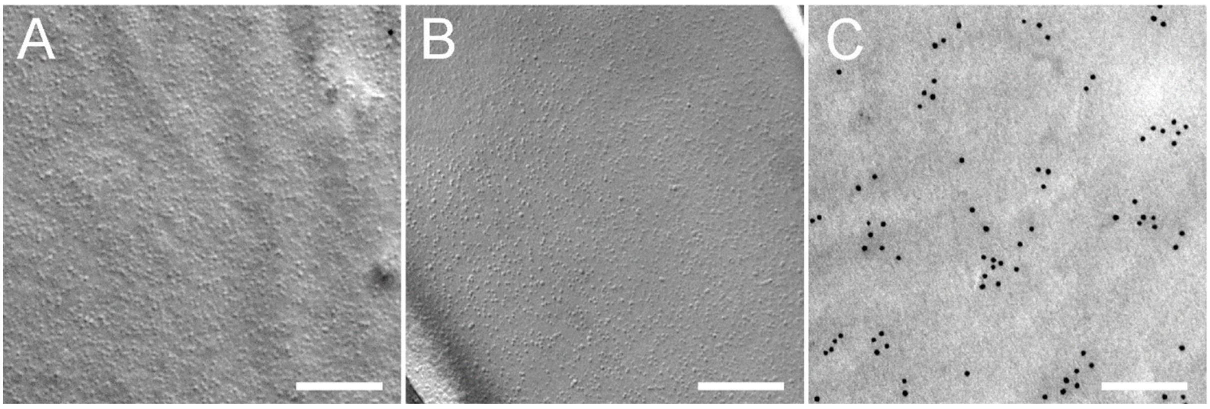


**SI Fig. S3: Negative and positive controls for SDS-FRL of *Nt*PINs in the cytosolic (P) side of tobacco cell PM fixed with high-pressure freezing. (A)** Negative control, preparations from wild-type exponential tobacco BY-2 cells immunostained with anti-GFP primary antibody and secondary antibody conjugated with colloidal gold (12 nm). **(B)** Negative control, preparations from induced XVE-*Nt*PIN11T-GFP immunostained only with secondary antibody conjugated with colloidal gold (12 nm). **(C)** Positive control showing PM-resident epitope. Preparations from wild-type exponential tobacco BY-2 cells immunostained with primary anti-phosphatidylinositol 4,5-bisphosphate (PIP2) antibody and secondary antibody conjugated with colloidal gold (12 nm).

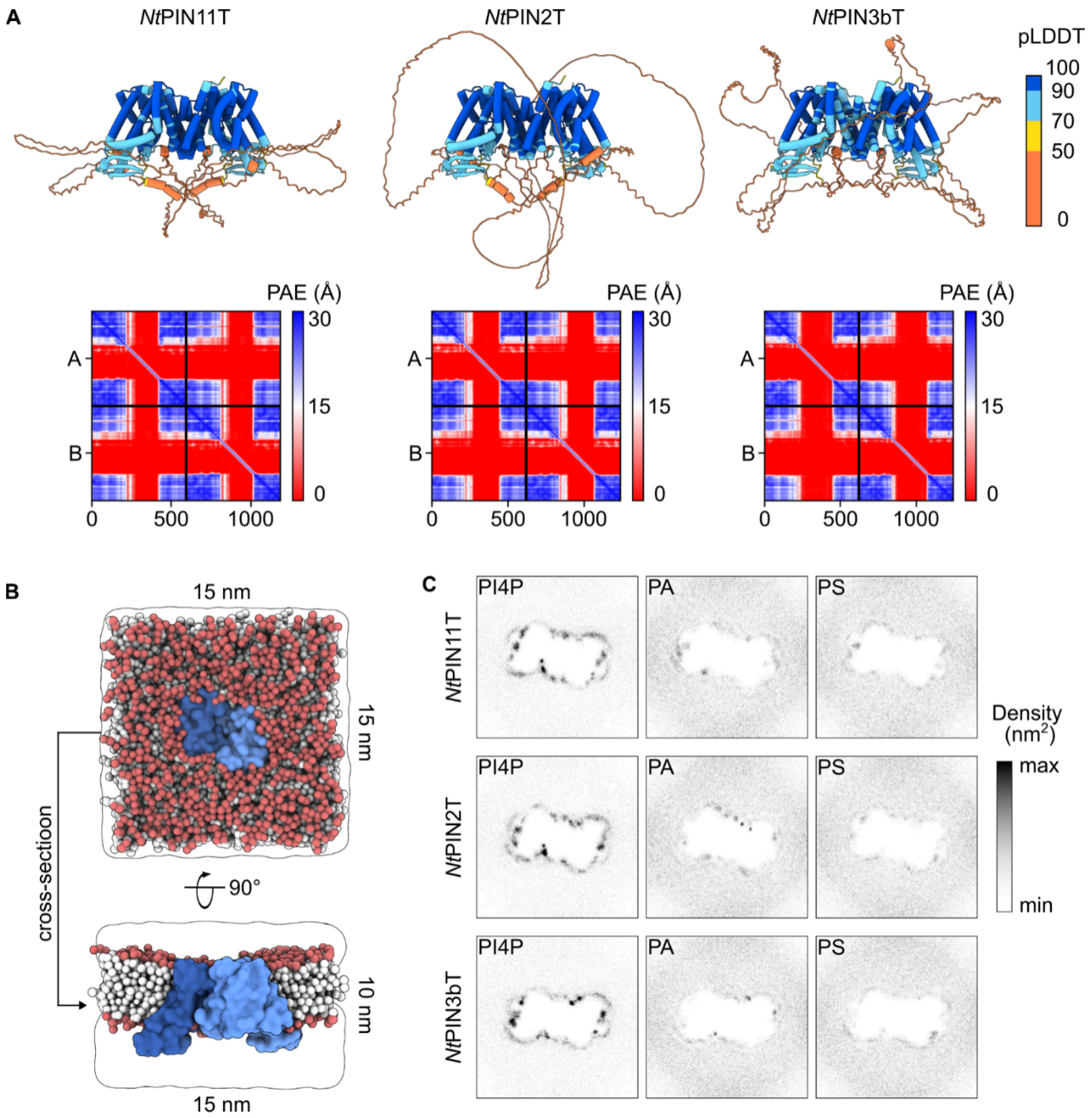


**SI Fig. S4: Lipid-protein interface of NtPINs. (A)** Full-length AlphaFold2 predictions of *Nt*PIN11T, *Nt*PIN2T and *Nt*PIN3bT dimers. For all *Nt*PINs the loop is predicted with the low pLDDT score. **(B)** Top and cross-section view on the simulation box containing the transmembrane region of *Nt*PIN11T dimer (shades of blue) surrounded by a model plasma membrane (lipid headgroups in red, lipid tails in white) and solvated in water and 0.15 M NaCl (white). Numbers indicate dimensions of the simulation box. **(C)** PI4P, PA and PS densities around transmembrane regions of *Nt*PIN11T, *Nt*PIN2T and *Nt*PIN3bT dimers. All *Nt*PINs dimers cluster primarily PI4P (and PI(4,5)P2, shown in Figure XY). No significant differences in clustering patterns among *Nt*PINs dimers were observed.

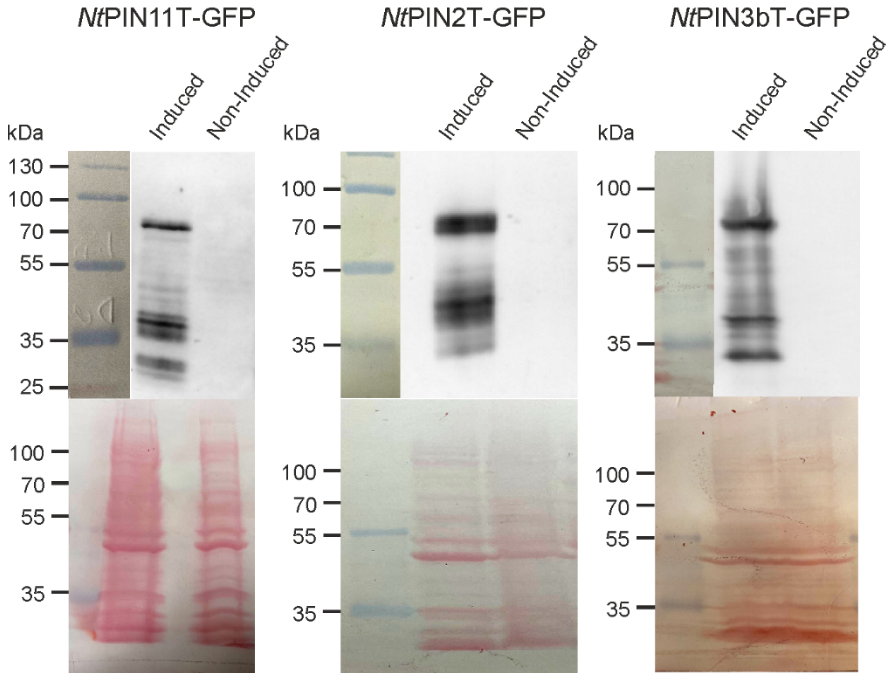


**SI Fig. S5: Immunodetection of GFP in CHAPS-solubilized membrane fractions used as input for co-IP.** Membrane fractions isolated from 2-day-old 1 µM β-estradiol induced and non-induced XVE-*Nt*PIN11T-GFP, *Nt*PIN2T-GFP, and *Nt*PIN3bT-GFP cells were separated by SDS-PAGE and immunoblots stained with anti-GFP primary antibody and secondary HRP-coupled secondary antibody followed by ECL chemiluminescence detection. The presence of proteins on membranes before immunodetection was tested by Ponceau S-staining (lower row).

**SI Table S1**: **List of candidate interactors identified by GFP-trap co-immunoprecipitation with *Nt*PIN11T-GFP analyzed by LC-MS$^2$.** 2-day-old XVE-*Nt*PIN11T-GFP cells were induced with 1 µM β-estradiol, PM fraction isolated, treated with CHAPS and analyzed with MS$^2$. Peptides were compared with the tobacco genome database (Edwards et al., 2017) using tblastn. The closest *Arabidopsis thaliana* homologs of individual proteins were found using blastp in TAIR10 protein list (Berardini et al., 2015). Shown are proteins that meet the following criteria: the average log2 intensity is ≥ 20 (32-fold higher abundance than the lowest intensity positively detected protein), the protein is identified in at least two of three biological repetitions, and the protein is strictly detected only in membrane fractions from induced cells, and not in membrane and cytosolic fractions from cells expressing free GFP. Interactors are color-coded as follows: cell wall proteins (yellow), cytosolic and membrane-bound enzymes (green), intracellular signaling and vesicle trafficking proteins (red). The abundance of bait (*Nt*PIN11T-GFP) detected by peptides corresponding to GFP was increased 1500-fold compared to the abundance of proteins with the lowest positively detected levels.

| ***Nicotiana tabacum* Gene ID** | **Average intensity (log2 scale)** | ***Arabidopsis thaliana* AGI locus code** | ***Arabidopsis thaliana* Protein name** |
|---|---|---|---|
| Nitab4.5_0006973g0030.1 | 22.87 | At5G03170 | **FLA11**; Fasciclin-like arabinogalactan protein 11 |
| Nitab4.5_0000073g0500.1 | 22.71 | At3G21790 | **UGT71B7**; UDP-Glycosyltransferase superfamily protein |
| Nitab4.5_0004320g0030.1 | 22.58 | At2G45470 | **FLA8**; Fasciclin-like arabinogalactan protein 8 |
| Nitab4.5_0000001g0430.1 | 22.01 | At1G64760 | **ZET**; Zerzaust, β-1,3 glucanase arabinogalactan protein |
| Nitab4.5_0000273g0080.1 | 21.97 | At5G05340 | **PRX52**; Peroxidase 52 |
| Nitab4.5_0001701g0170.1 | 21.94 | At4G12730 | **FLA2**; Fasciclin-like arabinogalactan protein 2 |
| Nitab4.5_0000396g0100.1 | 21.40 | At5G65720 | **NIFS1**; Nitrogen fixation S-like 1 protein |
| Nitab4.5_0003730g0080.1 | 21.15 | At4G33060 | **PPIase**; Cyclophilin-like peptidyl-prolyl cis-trans isomerase |
| Nitab4.5_0003321g0020.1 | 21.05 | At1G02130 | **RabD2a**; GTPase from Rab family |
| Nitab4.5_0000045g0200.1 | 20.95 | At2G32720 | **CB5-B**; Cytochrome B5 isoform B protein |
| Nitab4.5_0000566g0240.1 | 20.29 | At4G26300 | **EMB1027**; Arginyl-tRNA synthetase class Ic |
| Nitab4.5_0004737g0040.1 | 20.28 | At2G39990 | **eIF3F**; Eukaryotic translation initiation factor 3, subunit F |
| Nitab4.5_0003141g0070.1 | 20.11 | At5G63870 | **PP7**; Serine/threonine phosphatase 7 |
| Nitab4.5_0012117g0020.1 | 20.11 | At1G52910 | **DUF1218**; Unknown protein, cell wall localization (SUBA5) |
| Nitab4.5_0000005g0010.1 | 20.08 | At4G04210 | **PUX4**; Plant UBX domain-containing protein 4 |

**SI Table S2**: **List of candidate interactors identified by GFP-trap co-immunoprecipitation with *Nt*PIN2T-GFP analyzed by LC-MS$^2$.** 2-day-old XVE-*Nt*PIN2T-GFP cells were induced with 1 µM β-estradiol, PM fraction isolated, treated with CHAPS and analyzed with MS$^2$. Peptides were compared with the tobacco genome database (Edwards et al., 2017) using tblastn. The closest *Arabidopsis thaliana* homologs of individual proteins were found using blastp in TAIR10 protein list (Berardini et al., 2015). Shown are proteins that meet the following criteria: the average log2 intensity is ≥ 20 (32-fold higher abundance than the lowest intensity positively detected protein), the protein is identified in at least two of three biological repetitions, and the protein is strictly detected only in membrane fractions from induced cells, and not in membrane and cytosolic fractions from cells expressing free GFP. Interactors are color-coded as follows: cell wall proteins (yellow), cytosolic and membrane-bound enzymes (green), intracellular signaling and vesicle trafficking proteins (red). The abundance of bait (*Nt*PIN2T-GFP) detected by peptides corresponding to GFP was increased 2500-fold compared to the abundance of proteins with the lowest positively detected levels.

| ***Nicotiana tabacum*** **Gene ID** | **Average intensity (log2 scale)** | ***Arabidopsis thaliana*** **AGI locus code** | ***Arabidopsis thaliana*** **Protein name** |
|---|---|---|---|
| Nitab4.5_0000848g0100.1 | 23.64 | At1G07910 | **RNL**; tRNA ligase |
| Nitab4.5_0012117g0020.1 | 23.20 | At1G52910 | **DUF1218**; Unknown protein, cell wall localization (SUBA5) |
| Nitab4.5_0000073g0500.1 | 23.14 | At3G21790 | **UGT71B7**; UDP-Glycosyltransferase superfamily protein |
| Nitab4.5_0010277g0010.1 | 21.97 | At3G58730 | **VHA-D**; Vacuolar-type $H^+$ -ATPase, subunit D |
| Nitab4.5_0006973g0030.1 | 21.36 | At5G03170 | **FLA11**; Fasciclin-like arabinogalactan protein 11 |
| Nitab4.5_0004737g0040.1 | 21.26 | At2G39990 | **eIF3F**; Eukaryotic translation initiation factor 3, subunit F |
| Nitab4.5_0006944g0020.1 | 21.06 | At1G28490 | **SYP61**; SYNTAXIN OF PLANTS 61, SNARE protein |
| Nitab4.5_0000052g0230.1 | 20.81 | At4G32760 | **TOL9**; TOM1-LIKE ubiquitin-binding ESCRT protein |
| Nitab4.5_0009955g0060.1 | 20.73 | At4G17530 | **RAB1C**; RAB GTPase homolog 1C |
| Nitab4.5_0000410g0380.1 | 20.68 | At1G52760 | **LysoPL**; Lysophospholipase, ascorbate peroxidase activity |
| Nitab4.5_0003335g0040.1 | 20.19 | At4G17070 | **PPIase**; Peptidyl-prolyl cis-trans isomerase-like3 |

**SI Table S3: List of candidate interactors identified by GFP-trap co-immunoprecipitation with *Nt*PIN3bT-GFP analyzed by LC-MS$^2$.** 2-day-old XVE-*Nt*PIN3bT-GFP cells were induced with 1 µM β-estradiol, PM fraction isolated, treated with CHAPS and analyzed with MS$^2$. Peptides were compared with the tobacco genome database (Edwards et al., 2017) using tblastn. The closest *Arabidopsis thaliana* homologs of individual proteins were found using blastp in TAIR10 protein list (Berardini et al., 2015). Shown are proteins that meet the following criteria: the average log2 intensity is ≥ 20 (32-fold higher abundance than the lowest intensity positively detected protein), the protein is identified in at least two of three biological repetitions, and the protein is strictly detected only in membrane fractions from induced cells, and not in membrane and cytosolic fractions from cells expressing free GFP. Interactors are color-coded as follows: cell wall proteins (yellow), cytosolic and membrane-bound enzymes (green), intracellular signaling and vesicle trafficking proteins (red), and other proteins including ribosomal proteins and some membrane carriers (no color). The abundance of bait (*Nt*PIN3bT-GFP) detected by peptides corresponding to GFP was increased 10500-fold compared to the abundance of proteins with the lowest positively detected levels.

| ***Nicotiana tabacum*** **Gene ID** | ***Arabidopsis thaliana*** **AGI locus code** | **Average intensity (log2 scale)** | ***Arabidopsis thaliana*** **Protein name** |
|---|---|---|---|
| Nitab4.5_0001364g0100.1 | At2G39050 | 26.59 | **EULS3**; Hydroxyproline-rich glycoprotein family protein |
| Nitab4.5_0000073g0500.1 | At3G21790 | 23.14 | **UGT71B7**; UDP-Glycosyltransferase superfamily protein |
| Nitab4.5_0003719g0020.1 | At1G44170 | 24.40 | **ALDH4**; Aldehyde dehydrogenase 4 |
| Nitab4.5_0000367g0040.1 | At1G44170 | 24.31 | **uL22y**; Ribosomal protein L22p/L17e family protein |
| Nitab4.5_0005869g0030.1 | At4G30610 | 23.81 | **BRS1**; BRI1 suppressor; extracellular serine carboxypeptidase |
| Nitab4.5_0012849g0010.1 | At5G57290 | 23.68 | **p3y**; 60S acidic ribosomal protein family |
| Nitab4.5_0001215g0120.1 | At3G08640 | 23.50 | **RER3**; Alphavirus core family protein |
| Nitab4.5_0003037g0060.1 | At2G38670 | 23.45 | **PECT1**; Ethanolamine-phosphate cytidyltransferase 1 |
| Nitab4.5_0011890g0010.1 | At5G24710 | 23.28 | **TWD40-2**; Transducin TPLATE complex protein |
| Nitab4.5_0003904g0040.1 | At2G40010 | 23.00 | **uL10z**;Ribosomal protein L10 family protein |
| Nitab4.5_0000034g0170.1 | At1G59900 | 22.41 | **IAR4**; Pyruvate dehydrogenase E1 component alpha subunit |
| Nitab4.5_0000973g0040.1 | At5G65930 | 22.34 | **KCBP**; Kinesin-like calmodulin-binding protein (ZWICHEL) |
| Nitab4.5_0002531g0060.1 | At2G36850 | 22.20 | **GSL8**; Glucan synthase-like family protein 8 |
| Nitab4.5_0000573g0120.1 | At3G11560 | 22.11 | **LETM1**; LETM1-like protein |
| Nitab4.5_0000505g0020.1 | At2G43400 | 22.09 | **ETFQO**; Ubiquinone oxidoreductase |
| Nitab4.5_0009593g0010.1 | At1G66400 | 21.96 | **CML23**; Calcium-binding protein |
| Nitab4.5_0000486g0060.1 | At1G15880 | 21.92 | **GOS11**; Golgi SNARE 11 protein |
| Nitab4.5_0003928g0050.1 | At1G65470 | 21.67 | **FAS1**; Chromatin assembly factor |
| Nitab4.5_0001565g0110.1 | At2G34357 | 21.60 | **ARSP**; ARM repeat ribosomal RNA processing protein |
| Nitab4.5_0011900g0010.1 | At4G01320 | 21.55 | **STE24**; Peptidase family M48 family protein |
| Nitab4.5_0018351g0010.1 | At1G64790 | 21.51 | **GCN1**; Protein kinase regulating transcription and translation |
| Nitab4.5_0003141g0070.1 | At5G63870 | 21.47 | **PP7**; Serine/threonine phosphatase 7 |
| Nitab4.5_0010734g0020.1 | At3G62020 | 21.44 | **GLP10**; Germin like protein 10 |
| Nitab4.5_0000895g0200.1 | At3G17410 | 21.42 | **CARK1**; Putative receptor-like cytoplasmic kinase 1 |
| Nitab4.5_0013181g0010.1 | At1G77590 | 21.38 | **LACS9**; Long chain acyl-CoA synthetase 9, lipid biosynthesis |
| Nitab4.5_0004556g0020.1 | AtT3G17970 | 21.38 | **TOC64**; Integral chloroplast outer membrane protein |
| Nitab4.5_0000968g0060.1 | At5G27520 | 21.33 | **PNC2**; Peroxisomal adenine nucleotide carrier 2 |
| Nitab4.5_0007790g0010.1 | At4G37830 | 21.32 | **GRX4**; Cytochrome c oxidase-like protein |
| Nitab4.5_0004071g0010.1 | At4G03550 | 21.08 | **GSL5**; Glucan synthase-like family protein 5 |
| Nitab4.5_0003730g0080.1 | At4G33060 | 21.03 | **PPIase**; Cyclophilin-like peptidyl-prolyl cis-trans isomerase |
| Nitab4.5_0000366g0060.1 | At5G55940 | 20.96 | **EMB2731**; Uncharacterized protein family |
| Nitab4.5_0000383g0190.1 | At3G50930 | 20.94 | **CM66**; Mitochondrial membrane AAA ATPase |
| Nitab4.5_0004737g0040.1 | At2G39990 | 20.80 | **eIF3F**; Eukaryotic translation initiation factor 3, subunit F |
| Nitab4.5_0003909g0020.1 | At1G43190 | 20.69 | **PTB3**; Polypyrimidine tract-binding protein 3, splicing |
| Nitab4.5_0000492g0070.1 | At1G63440 | 20.43 | **HMA5**; Plasma membrane P-type ATPase, primary transporter |
| Nitab4.5_0001381g0050.1 | At1G60170 | 20.38 | **PRP31**; Splicing factor |
| Nitab4.5_0006973g0030.1 | At5G03170 | 20.25 | **FLA11**; Fasciclin-like arabinogalactan protein 11 |
| Nitab4.5_0001241g0030.1 | At4G27480 | 20.08 | **GT14**; Glycosyltransferase family14 |

**SI Table S4**: **List of fold-change candidate interactors identified by GFP-trap co-immunoprecipitation with *Nt*PINs analyzed by LC-MS$^2$.** 2-day-old XVE-*Nt*PIN2T-GFP, XVE-*Nt*PIN3bT-GFP and XVE-*Nt*PIN11T-GFP cells were induced with 1 µM β-estradiol, PM fraction isolated, treated with CHAPS and analyzed with MS$^2$. Peptides were compared with the tobacco genome database (Edwards et al., 2017) using tblastn. The closest *Arabidopsis thaliana* homologs of individual proteins were found using blastp in TAIR10 protein list (Berardini et al., 2015). Individual candidates were manually filtered against profiles from non-induced cells and cells expressing free GFP. Three biological repetitions were performed, identified candidates were present in at least two of them. Shown are only manually selected PM-related proteins that have more than 10-fold enrinchment in the identified peptides. Interactors are color-coded as follows: PM-related enzymes (blue), endocytosis-related proteins (green), and vesicle trafficking proteins (red).

| ***Nicotiana tabacum*** **Gene ID** | ***Arabidopsis thaliana*** **AGI locus code** | **Average fold change** | ***Arabidopsis thaliana*** **Protein name** |
|---|---|---|---|
| Nitab4.5_0000055g0200.1 | At4G30190 | 20 (*Nt*PIN2T x free GFP)<br>31 (*Nt*PIN3bT x non-induced)<br>177 (*Nt*PIN3bT x free GFP) | **AHA2**; Plasma membrane $H^+$ ATPase 2 |
| Nitab4.5_0007642g0010.1 | At5G62670 | 11 (*Nt*PIN3bT x non-induced)<br>39 (*Nt*PIN3bT x free GFP) | **AHA11**; Plasma membrane $H^+$ ATPase 11 |
| Nitab4.5_0000336g0120.1 | At1G49340 | 22 (*Nt*PIN3bT x non-induced) | **PI4K ALPHA**; Phosphatidylinositol 4-kinase alpha |
| Nitab4.5_0001127g0070.1 | At2G42010 | 11 (*Nt*PIN2T x non-induced)<br>39 (*Nt*PIN3bT x free GFP) | **PLD BETA1**; Phospholipase D BETA 1 |
| Nitab4.5_0001402g0010.1 | At2G42910 | 114 (*Nt*PIN3bT x free GFP) | **PRS4**; Phosphoribosyltransferase family protein |
| Nitab4.5_0004506g0080.1 | At5G42080 | 11 (*Nt*PIN2T x free GFP)<br>29 (*Nt*PIN3bT x non-induced)<br>95 (*Nt*PIN3bT x free GFP) | **DRP1A**; Dynamin-related protein |
| Nitab4.5_0006873g0010.1 | At3G60190 | 46 (*Nt*PIN2T x free GFP)<br>934 (*Nt*PIN3bT x free GFP) | **DRP1E**; Dynamin-related protein |
| Nitab4.5_0001057g0100.1 | At1G59610 | 58 (*Nt*PIN3bT x free GFP) | **DRP2B**; Dynamin-related protein |
| Nitab4.5_0001773g0030.1 | At4G33650 | 56 (*Nt*PIN3bT x free GFP) | **DRP3A**; Dynamin-related protein |
| Nitab4.5_0006114g0010.1 | At2G07360 | 36 (*Nt*PIN3bT x non-induced) | **TASH3**; TPLATE-associated SH3 domain protein |
| Nitab4.5_0014777g0010.1 | At1G43890 | 65 (*Nt*PIN2T x non-induced) | **RAB18**; Ras-related small GTPase 18.3 |
| Nitab4.5_0001417g0080.1 | At5G67560 | 20 (*Nt*PIN3bT x free GFP) | **ARL8B**; Small GTPase Rab-type protein |
| Nitab4.5_0001486g0050.1 | At4G09720 | 14 (*Nt*PIN3bT x free GFP) | **RABG3A**; Small GTPase Rab-type protein |
| Nitab4.5_0002056g0020.1 | At1G04760 | 17 (*Nt*PIN3bT x free GFP) | **VAMP726**; Synaptobrevin-like protein family |
| Nitab4.5_0003114g0050.1 | At1G61250 | 35 (*Nt*PIN3bT x free GFP) | **SCAMP1**; Secretory carrier membrane protein |

#### SI References

Berardini, T. Z., Reiser, L., Li, D., Mezheritsky, Y., Muller, R., Strait, E., & Huala, E. (2015). The arabidopsis information resource: Making and mining the “gold standard” annotated reference plant genome. *Genesis*, *53*(8), 474-485.

Edwards, K. D., Fernandez-Pozo, N., Drake-Stowe, K., Humphry, M., Evans, A. D., Bombarely, A., Allen, F., Hurst, R., White, B., Kernodle, S. P., Bromley, J. R., Sanchez-Tamburrino, J. P., Lewis, R. S., & Mueller, L. A. (2017). A reference genome for Nicotiana tabacum enables map-based cloning of homeologous loci implicated in nitrogen utilization efficiency. *BMC Genomics*, *18*(1), 1-14.
